# The MexMAGIC population reveals the genetic architecture of clinal trait variation in Mexican native maize

**DOI:** 10.64898/2026.01.17.700095

**Authors:** Sergio Perez-Limón, Ana Laura Alonso-Nieves, M. Rosario Ramírez-Flores, G. Carolina Cintora-Martínez, M. Nancy Salazar-Vidal, Jessica Carcaño-Macías, Melanie G. Perryman, Oliver H. S. Paulson, Jagdeep S. Sidhu, Peng Yu, Victor Llaca, Kevin A. Fengler, Forrest Li, Daniel E. Runcie, Jeffrey Ross-Ibarra, C. Stewart Gillmor, Rubén Rellán-Álvarez, Ruairidh J. H. Sawers

**Affiliations:** Department of Plant Science, The Pennsylvania State University, University Park, PA 16802, USA; Unidad de Genómica Avanzada, Cinvestav, Irapuato, 36824, Guanajuato, México; Departamento de Fitomejoramiento, Universidad Autónoma Agraria Antonio Narro, Calzada Antonio Narro 1923, Colonia Buenavista, Saltillo, Coahuila, 25315, México; Departamento de Biotecnología y Bioquímica, Centro de Investigación y de Estudios Avanzados (CINVESTAV-IPN), Irapuato, Guanajuato, 36821, México; Division of Plant Science and Technology, University of Missouri, Columbia, MO 65211, USA; Department of Plant Genetics, TUM School of Life Sciences, Technical University of Munich, Germany; Corteva Agriscience, Johnston, IA, USA; Department of Evolution and Ecology, University of California, Davis, CA, USA 95616; Department of Plant Sciences, University of California, Davis, CA, USA 95616; Department of Molecular and Structural Biochemistry, N.C. Plant Sciences Initiative, North Carolina State University, Raleigh, NC 27695

## Abstract

Defining the genetic basis of local adaptation is fundamental to evolutionary biology and crop improvement. Theory predicts that when selective pressures track differences in the environment, a cline will be established. Such clines might be exploited to uncover adaptive variation by association of alleles with environmental stressors. However, monotonic phenotypic change over a cline is not necessarily mirrored by adaptive genetic variants. Furthermore, population structure can complicate the interpretation of genotype-environment association. To test the assumptions of genotype-environment association in a crop species, we developed a multi-parent advanced generation inter-cross (MAGIC) population using eight Mexican native maize varieties sourced from distinct agroecological zones. We mapped two clinal traits (tassel branching and flowering time) differing in genetic architecture. Variation in tassel branch number was dominated by a single QTL with allele effects that aligned well with a negative elevational cline. In contrast, we mapped 11 flowering time QTL with allele effects that were not consistently correlated with any one source environmental factor and distinct loci donated by highland and lowland early maturing varieties. Our observations support the theoretical result that genotype-environment association will be strongest under simple genetic architecture, although identification of adaptive alleles may still be confounded by population structure.

**Plain Language:** Nine thousand years of careful selection and cultivation by indigenous farmers has generated a rich diversity of native Mexican maize (corn) varieties, grown from sea level to high mountains, and from jungle to semidesert. By crossing native varieties adapted to different locations, we can uncover important genetic variants conferring tolerance to environmental stressors.

## INTRODUCTION

A cline is a pattern of monotonic change in a biological character over an environmental gradient (Huxley, 1938; Savolainen *et al*., 2013; Lotterhos, 2023). Population genetic theory predicts that clinal variation will arise across continuous populations when selective pressures track differences in the environment (Huxley, 1938; Haldane, 1948; Fisher, 1950). Many empirical examples of clinal variation have been identified over both geographic axes (*e.g.* latitude and elevation) and gradients of specific environmental factors (*e.g.* temperature, precipitation). While not all phenotypic variation is adaptive, clines can be exploited to generate hypotheses regarding the adaptive value of a trait with respect to a given environmental pressure (Stinchcombe *et al*., 2004; Koski & Ashman, 2015). Such inferences can be greatly strengthened by assessing repeatability across parallel clines, although this itself makes assumptions about contingency and reproducibility in trait selection.

With the availability of large scale genomic datasets linked to collections of geo-referenced biological diversity, cline theory can be used to support the direct identification of adaptive alleles through genotype-environment association (GEA; Lasky *et al*., 2023). GEA is trait agnostic and relies only on correlation between allele frequency and source environment. The genome can be scanned for GEAs in an environmental Genome Wide Association (GWA), providing an attractive opportunity to mine broad diversity for useful variation without the need for logistically challenging and costly phenotypic evaluation (*e.g.* Key et al., 2018; Mao *et al*., 2019; Li *et al*., 2025). Although environmental GWA has great potential, it rests on assumptions that may, or may not, hold for a given study system.

Isolation by distance along a cline will almost inevitably result in limitations to geneflow that can establish population structure aligned with, and thereby confounding, the genetic signals underlying adaptation (Ahrens *et al*., 2021; Lasky *et al*., 2023). As such, control of population structure in an environmental GWA, while necessary, risks eliminating the very signals that are the focus of the study. More fundamentally, GEA assumes that gradual monotonic change in adaptive traits over an environmental cline is reflected by similar behavior in allele frequency at the causative loci. The validity of this assumption will be greatly impacted by the genetic architecture of underlying adaptive traits, *i*.*e*. the number of causative loci, the size and specificity of their effects, and their interactions with each other and with the environment. As genetic architecture becomes more complex, the effect of any given allele may be small, and indeed even neutral over sections of the landscape. In addition, individuals equally adapted to a particular locality might harbor distinct, but functionally equivalent, complements of adaptive alleles. Under such a scenario, the frequency of any given allele may be only weakly correlated with the environment even if the emergent patterns of local adaptation at the whole genotype level are strong.

Maize was domesticated from Balsas teosinte (*Z. mays* ssp. *parviglumis*; Matsuoka *et al*., 2002) by indigenous peoples at least 9,000 years ago in the basin of the Balsas River in Mexico (Matsuoka *et al*., 2002; Piperno *et al*., 2007). Mexico remains a center of maize diversity, with 59 different documented native varieties (Sanchez G. *et al*., 2000). Although each variety has its own character and associated culinary and cultural importance (Louette *et al*., 1997; Ortega-Paczka, 2003; Cleveland & Soleri, 2007; Mercer & Perales, 2019), overlapping distribution and inter-varietal hybridization mean that Mexican native maize varieties can be considered a continuous, if highly structured, population (Ortega-Paczka, 2003; Campbell *et al*., 2025). Mexican native maize is cultivated from the tropical jungles of Chiapas to the highland semideserts of Chihuahua, from sea level to ∼3,400 m.a.s.l. (Ruiz Corral et al. 2008; www.biodiversidad.gob.mx). Previous characterization has documented morphological clines and local adaptation in Mexican native maize (Wellhausen *et al*., 1951; Eagles & Lothrop, 1994; Sanchez G. *et al*., 2000; Perez-Limón *et al*., 2022; Mercer *et al*., 2008; Romero Navarro *et al*., 2017; Janzen *et al*., 2022). Coupled with the substantial genomic resources available for maize, Mexican native maize represents a compelling system for the study of local adaptation.

The genetic basis of local adaptation in Mexican maize has been extensively investigated through comparative genomics, and both phenotypic and environmental GWA (Romero Navarro *et al*., 2017; Wang *et al*., 2021; Romero Navarro *et al*., 2017; Li *et al*., 2025). To complement such work and address the difficulty presented by strong environmentally-correlated population structure, we previously developed a biparental linkage mapping population using a single highland Mexican maize variety and a reference inbred line (Janzen *et al*., 2022; Perez-Limón *et al*., 2022). Here, we extend this approach with the development a multi-parent advanced generation inter-cross population (MexMAGIC) population that samples broader diversity and breaks population structure while retaining the power of linkage mapping (Scott *et al*., 2020). Significantly, as the MexMAGIC population captures more than one Mexican maize haplotype, allele effects can be estimated across the sampled environmental range and commonalities and differences assessed among the founder haplotypes. Here, we present the mapping of traits differing in complexity of genetic architecture (kernel pigmentation, tassel branching and flowering time). Trait data was collected over two seasons in a single common garden environment in Mexico. We discuss our results in the context of the genetic architecture of local adaptation and their implications for GEA approaches.

## RESULTS

### MexMAGIC founders were sourced from across the range of Mexican maize cultivation

To characterize the genetic architecture of trait variation and local adaptation, we generated a multi-parent advanced generation inter-cross (MAGIC) population (Scott *et al*., 2020) using Mexican native maize, sourced to capture eight founder haplotypes from across the breadth of the Mexican environment (MexMAGIC; hereafter, we refer to the founder varieties by three letter identifiers given in Table 1, Table S1). The eight founders were selected to sample different previously-described climatic-adaptation groups (Eagles & Lothrop, 1994; Ruiz Corral *et al*., 2008) and, when compared to a previously characterized broad sampling of Mexican native maize accessions (Romero Navarro *et al*., 2017), they showed good coverage of both geographical (Fig. 1A) and environmental (Fig. 1B; S1-4; Table S2, 3) range. For pragmatic reasons, we used inbred (S_6_) lines (Chia *et al*., 2012; Venkatesh *et al*., 2015) as founders where available, although it remained necessary to source five of the eight founders from outbred open pollinated varieties (OPVs) to sample the environments of interest.

**Table 1.**
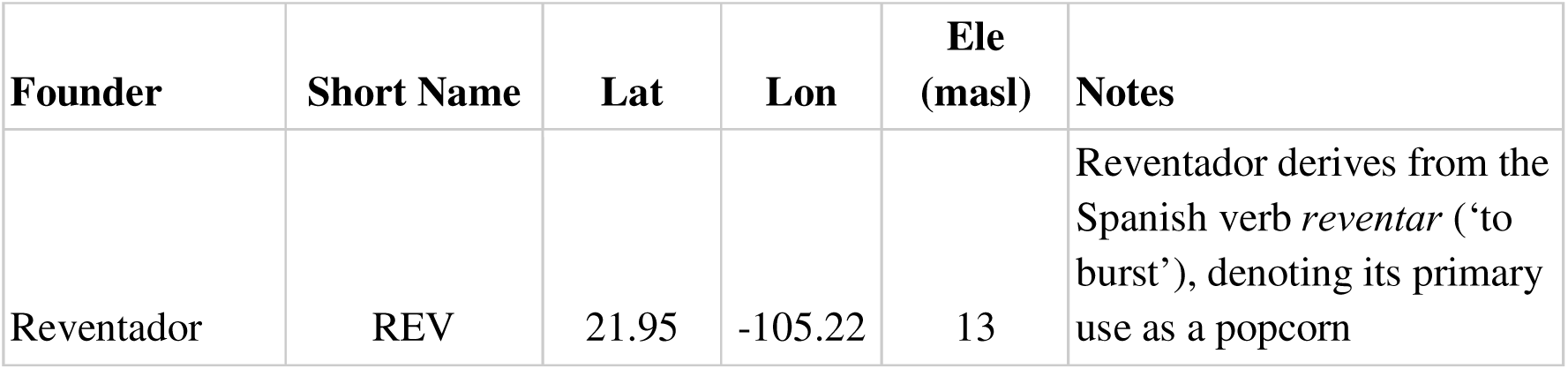
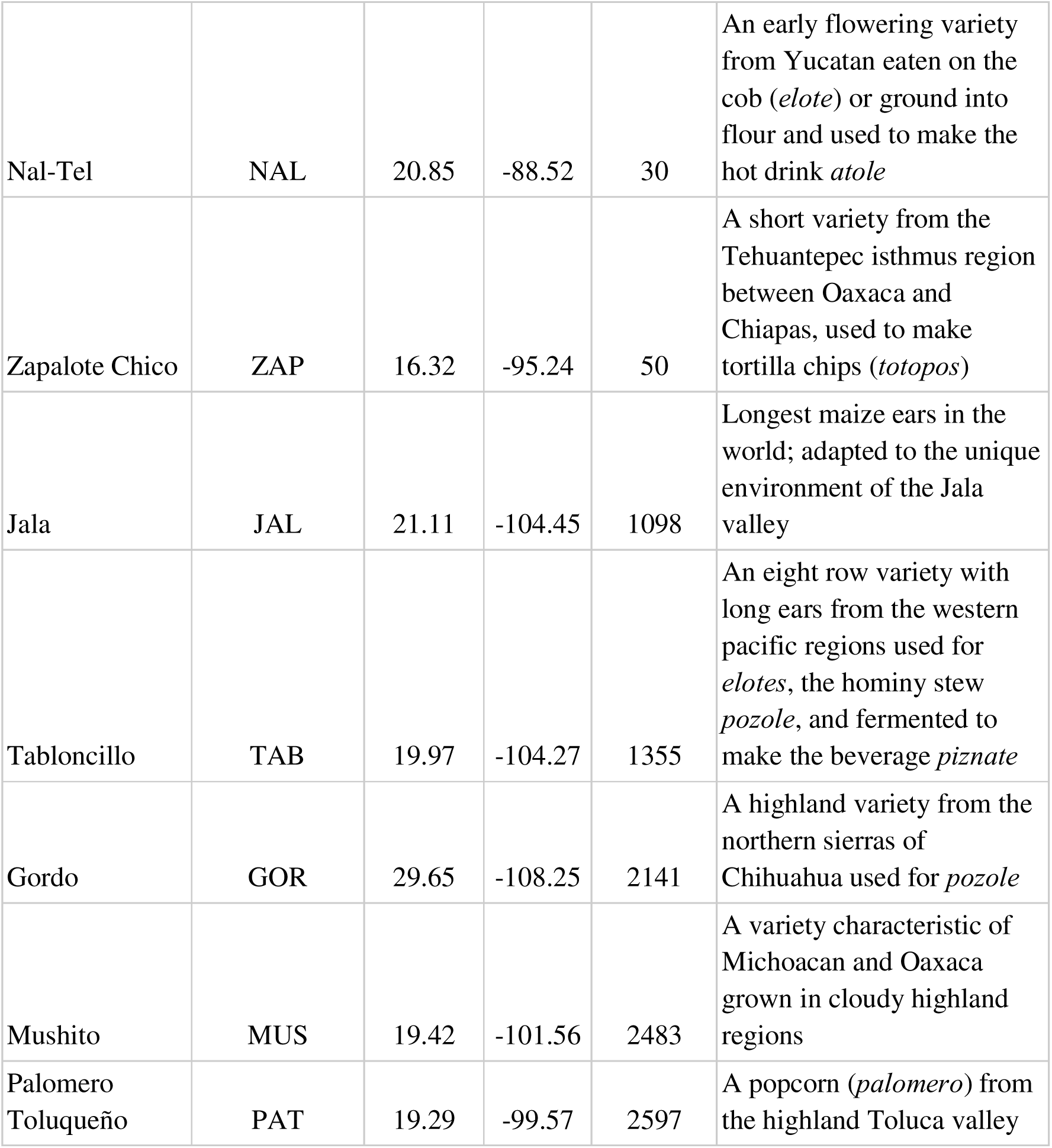
MexMAGIC founders

| Founder | Short Name | Lat | Lon | Ele<br>(masl) | Notes |
| --- | --- | --- | --- | --- | --- |
| Reventador | REV | 21.95 | -105.22 | 13 | Reventador derives from the Spanish verb *reventar* ('to burst'), denoting its primary use as a popcorn |
| Nal-Tel | NAL | 20.85 | -88.52 | 30 | An early flowering variety from Yucatan eaten on the cob (*elote*) or ground into flour and used to make the hot drink *atole* |
| Zapalote Chico | ZAP | 16.32 | -95.24 | 50 | A short variety from the Tehuantepec isthmus region between Oaxaca and Chiapas, used to make tortilla chips (*totopos*) |
| Jala | JAL | 21.11 | -104.45 | 1098 | Longest maize ears in the world; adapted to the unique environment of the Jala valley |
| Tabloncillo | TAB | 19.97 | -104.27 | 1355 | An eight row variety with long ears from the western pacific regions used for *elotes*, the hominy stew *pozole*, and fermented to make the beverage *piznate* |
| Gordo | GOR | 29.65 | -108.25 | 2141 | A highland variety from the northern sierras of Chihuahua used for *pozole* |
| Mushito | MUS | 19.42 | -101.56 | 2483 | A variety characteristic of Michoacan and Oaxaca grown in cloudy highland regions |
| Palomero Toluqueño | PAT | 19.29 | -99.57 | 2597 | A popcorn (*palomero*) from the highland Toluca valley |

**Figure 1.**
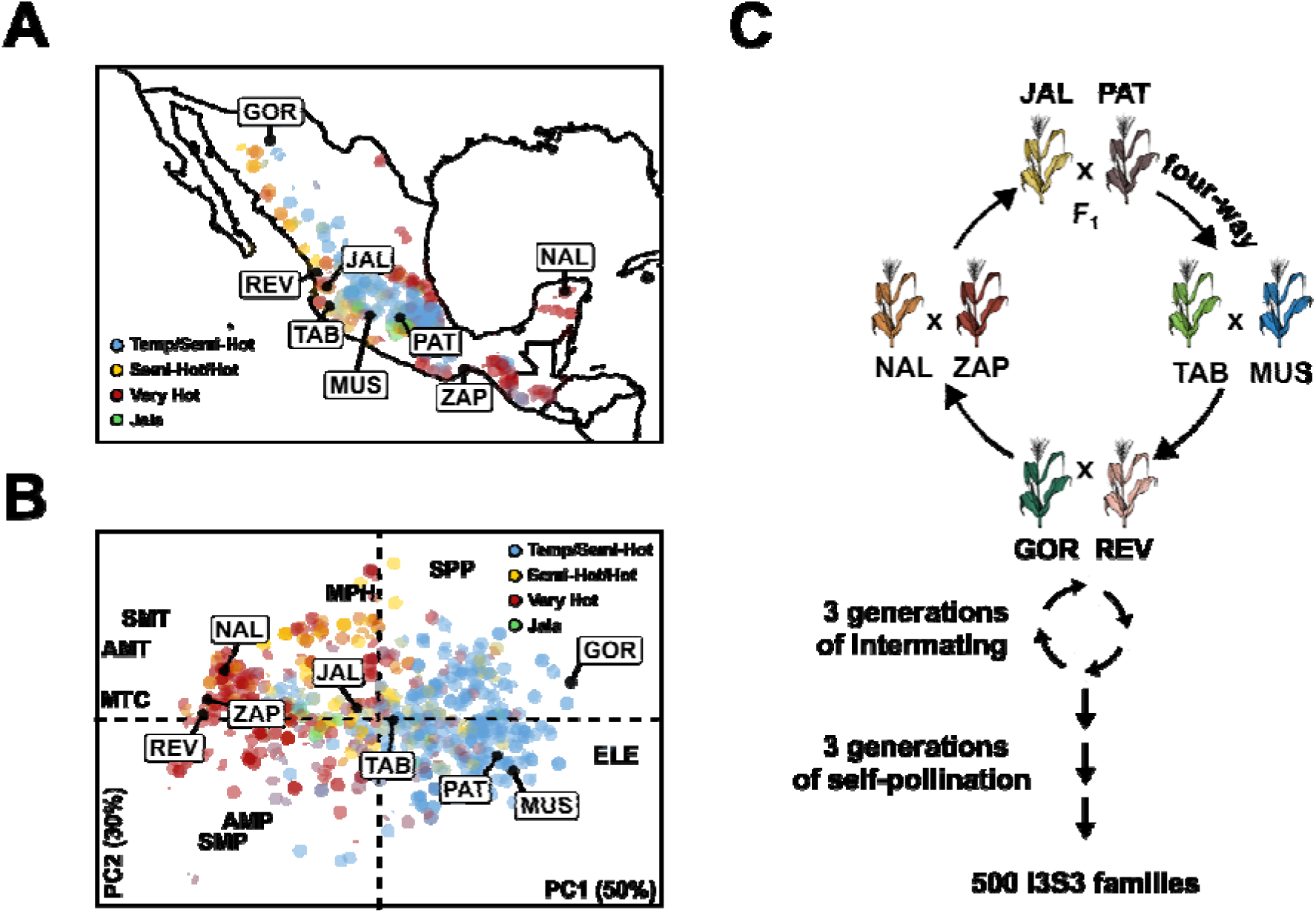
Founder selection and design of the MexMAGIC population. **A.** Source of the eight founder varieties (boxed three letter identifiers) in the context of a broader Mexican diversity panel (Romero Navarro *et al*., 2017) labeled by previously assigned agroecological environment (Eagles & Lothrop, 1994; Ruiz Corral *et al*., 2008). **B.** Environmental principal component (PC) analysis of MexMAGIC founders (boxed three letter identifiers) and Mexican diversity panel based on elevation (ELE), seasonal photoperiod (SPP), annual mean temperature (AMT), seasonal mean temperature (SMT), annual mean precipitation (AMP), seasonal mean precipitation (SMP), mean temperature of the coldest month (MTC) and mean soil pH (MPH). Biplot shows PC1 (explaining 50% variance) and PC2 (explaining 30% variance). Position of environmental descriptor labels indicates relative importance and sign of loading. Variety points colored as panel A. **C.** Crossing scheme for generation of MexMAGIC from 8 founder parents.

Consequently, we restricted the second generation in our breeding scheme to four single F_1_ individuals, each derived from a distinct hybrid cross, ensuring that each founder donated only a single haplotype to the final population (Fig. 1C, S5-7). After subsequent rounds of intermating (see Materials and Methods), we generated > 500 MexMAGIC partially inbred (S_3_) families. Here, we present genetic and phenotypic characterization of a first cohort of 200 families. For phenotypic evaluation, families were test-crossed to the inbred line NC358 and evaluated over two years in a mid-elevation Mexican field site.

### The genomes of the MexMAGIC families are mosaics of the founder parents

To determine the eight MexMAGIC founder haplotypes, we generated ∼30x whole genome sequence (WGS) data for the four founding F_1_ individuals, two of the three inbred founders (TAB, REV), the OPV JAL parent, and related individuals sampled from the founder OPVs (Table S1). We had previously generated WGS data for the PAT founder (Gonzalez-Segovia *et al*., 2019). Similarly, further WGS data was available for the three inbred founders (TAB, REV, ZAP) from an earlier maize HapMap study (Chia *et al*., 2012; Venkatesh *et al*., 2015). While working on the MexMAGIC, we also generated complete genome assemblies for TAB and ZAP inbreds, along with a haploid assembly for an individual sampled from same PAT OPV accession used in the MexMAGIC population (www.maizegdb.org/HiLo_project). For each F_1_ genotype, we used a triplet approach to phase heterozygous sites with respect to the founder parents and determined the founder haplotypes (Fig. S8A). Overall, across the eight founder haplotypes, we identified 14×10^6^ polymorphic sites that were assembled into a haplotype graph for subsequent analysis of the derived MexMAGIC families (Bradbury *et al*., 2022).

The 200 MexMAGIC families (Fig. S8B) were genotyped using a 50k Single Nucleotide Polymorphism (SNP) array. After filtering, we obtained ∼10k polymorphic SNPs (Table S4). Principal coordinate (PCo) analysis showed the MexMAGIC families to be broadly scattered across the genetic space defined by the eight founders (Fig. S8C; Table S5). There were several families that showed outlying loading on PCo1 (11 greater than 2SDs above the mean) that was distinct from other families and the founders (Fig. S8C, 9), although when analyzed at the individual chromosome level, loading on PCo1 fell within the range defined by the founders for all families (Fig. S10). Given that phenotypic data was available, we chose to retain all 200 MexMAGIC for subsequent analysis. PCo2 nicely separated the founders by source elevation with good coverage of MexMAGIC families over the range (Fig. S8C,D; Table S5). To increase the mapping resolution and the SNP density in the MexMAGIC families, the SNPs identified with the genotyping array were used as a scaffold to impute the founder WGS SNPs into the families using the PHG v1.0 imputation pipeline. After the imputation process and filtering for missing data, 8 million polymorphic SNPs were imputed per family. To improve computational efficiency and mapping quality, markers with the same segregation pattern were filtered out, and a final set of 17,030 markers was obtained. The SNP-level data of the founders and MexMAGIC families was used to generate haplotype-based genome-wide assignment of founder genotype probability for each family (Fig. S8E). We also used hard calls of founder ancestry across the genome to infer gene-level allele origin as an aid to subsequent candidate-gene based analyses (Table S6).

### Major kernel color loci segregate in the MexMAGIC population

To assess the utility of the MexMAGIC population in QTL mapping, we characterized two highly heritable kernel pigment traits associated with well-defined large effect loci (Chen *et al*., 2024). We first mapped the accumulation of red flavonoid-derived phlobaphenes in the maternal pericarp tissue, known to be regulated by the *myb*-like transcription factor *P1* (Grotewold *et al*., 1994). In the MexMAGIC, red kernel pigmentation was unambiguously donated by the REV parent - red pigmentation was only seen to segregate in the REV OPV and the derived S6 inbred used as a founder (Fig. 2A). We then characterized yellow carotenoid pigments in the endosperm, known to be catalyzed by a phytoene synthase encoded by the *Y1* gene (Buckner *et al*., 1996; Palaisa *et al*., 2004). Yellow pigmentation segregated in NAL, PAT and MUS OPVs (Fig. 2A). The JAL x PAT F1 ear did not segregate yellow kernels, indicating that PAT did not donate an active *Y1* allele to the population.

**Figure 2.**
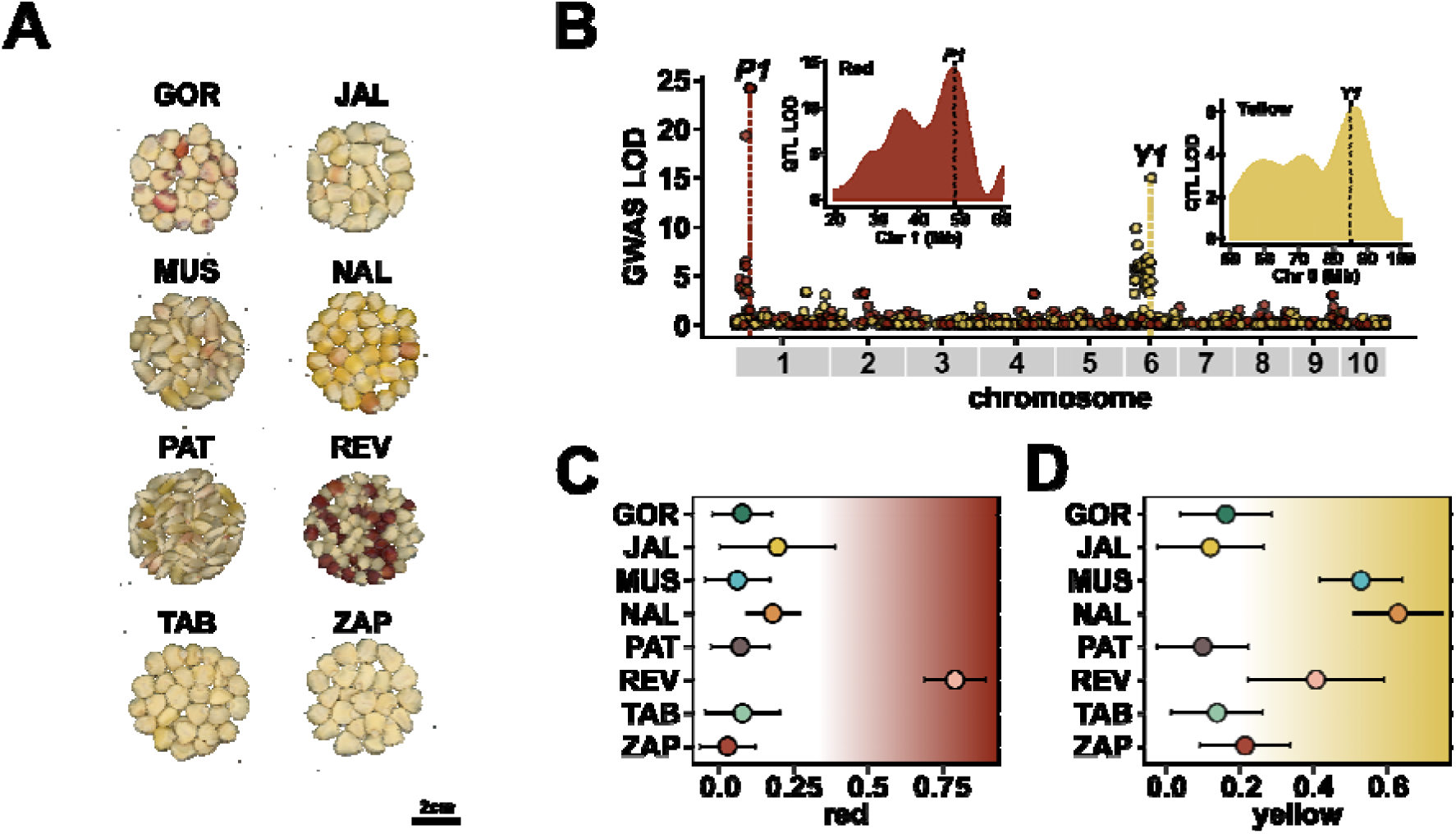
Classical loci are associated with variation in pericarp and aleurone pigmentation in the MexMAGIC. **A.** Kernels of the eight MexMAGIC founder accessions. The inbred founders REV, TAB and ZAP are represented here by related open-pollinated accessions. **B.** Main plot shows GWAS support (LOD) for association with red pericarp (red points) or yellow aleurone (yellow points) pigmentation across the ten maize chromosomes. All SNPs with LOD > 3 are shown. Only a reduced random sampling of lower-supported SNPs is shown. The physical position of the candidate genes *P1* and *Y1* is indicated by a dashed vertical line. Inset plots show smoothed LOD support for 8-allele QTL models for red and yellow pigmentation around *P1* and *Y1*, respectively. **C., D.** Estimated founder allele-effects at markers linked to *P1* and *Y1* for red and yellow pigmentation, respectively. Pigmentation (red or yellow) was scored per family as a binary trait, with 0 as not pigmented, and 1 as pigmented. Points show BLUPs of QTL effects for each founder. Bars show **±** Standard Error.

Unfortunately, pigmentation in the F1 ears TAB x MUS and NAL x ZAP was not recorded, although three of the four four-way ears segregated yellow kernels, indicating that both MUS and NAL had donated *Y1* alleles (Fig. 2C, S5). These segregation patterns provided a clear expectation of not only the loci but also the parental allele effects anticipated to underlie kernel pigmentation in the MexMAGIC.

We scored red and yellow kernel pigmentation as binary traits (presence/absence) across 196 MexMAGIC families (Table S7). Red pigmented families were removed from the yellow analysis due to the ambiguity of scoring pigmentation in the aleurone underlying the red pericarp. Families that showed pigment segregation (in the case of red, at the ear level due to segregation in maternal genotype; in the case of yellow, at the kernel level, due to segregation of the endosperm genotype) were scored as pigmented. The frequency of red and yellow families in the population corresponded to our expectation of 0.125 and 0.375 respectively (χ^2^ = 1.23, df = 1, p-value = 0.27; χ^2^ = 0.18, df = 1, p-value = 0.67). Running SNP based GWAS, we identified lead hits at 11 kb from *P1* (SNP 1_48412077) and 40 kb from *Y1* (SNP 6_85019028), for red and yellow pigmentation, respectively (Fig. 2B, Table S8). We then ran QTL models fitting the eight parental alleles, again recovering major peaks closely linked to *P1* and *Y1* candidate loci (Fig. 2B). Estimation of parental allele effects identified REV as the donor of the red pigmentation, and MUS and NAL as the strongest yellow donors, in line with our expectation (Fig. 2C, D). Overall, analysis of kernel pigmentation provided validation of genotyping, the genetic map and inference of founder ancestry across the MEMA family genomes.

### The MexMAGIC founders capture an elevational cline in tassel branch number

Having mapped kernel pigmentation in the MexMAGIC, we next sought to characterize the genetic architecture underlying a well-established elevational cline in tassel (male inflorescence) branch number (TBN). Native Mexican highland maize produces large tassels with few or no lateral branches, while the tassels of lowland varieties are highly branched (up to 40 branches), resembling the maize ancestor *Zea mays* ssp. *parviglumis* (Wellhausen et al., 1951; Eagles & Lothrop, 1994; Sanchez G. et al., 2000; Perez-Limón et al., 2022). A prior study of phenotypic and genetic diversity in native maize varieties (HiLo panel; Fig. 3B; S13) reported an excess of Qst/Fst associated with TBN between lowland and highland Mexican maize, consistent with locally acting selection (Janzen *et al*., 2022). We characterized the population structure of this previously studied HiLo panel and found a strong correlation with both elevation and TBN (Fig. S14,15), illustrating a fundamental challenge to identification of loci using GEA. In this context, the reduced population structure in the MexMAGIC presents an advantage. We evaluated the eight MexMAGIC founder varieties and found that they followed the TBN cline with respect to source elevation (Fig. 3A), indicating the population to be suitable for investigation of the underlying genetic architecture. Specifically, we aimed to use the MexMAGIC lines to evaluate evidence for 1) clinal allelic series and 2) bias in the directionality of genome-wide allele effects with respect to source elevation.

**Figure 3.**
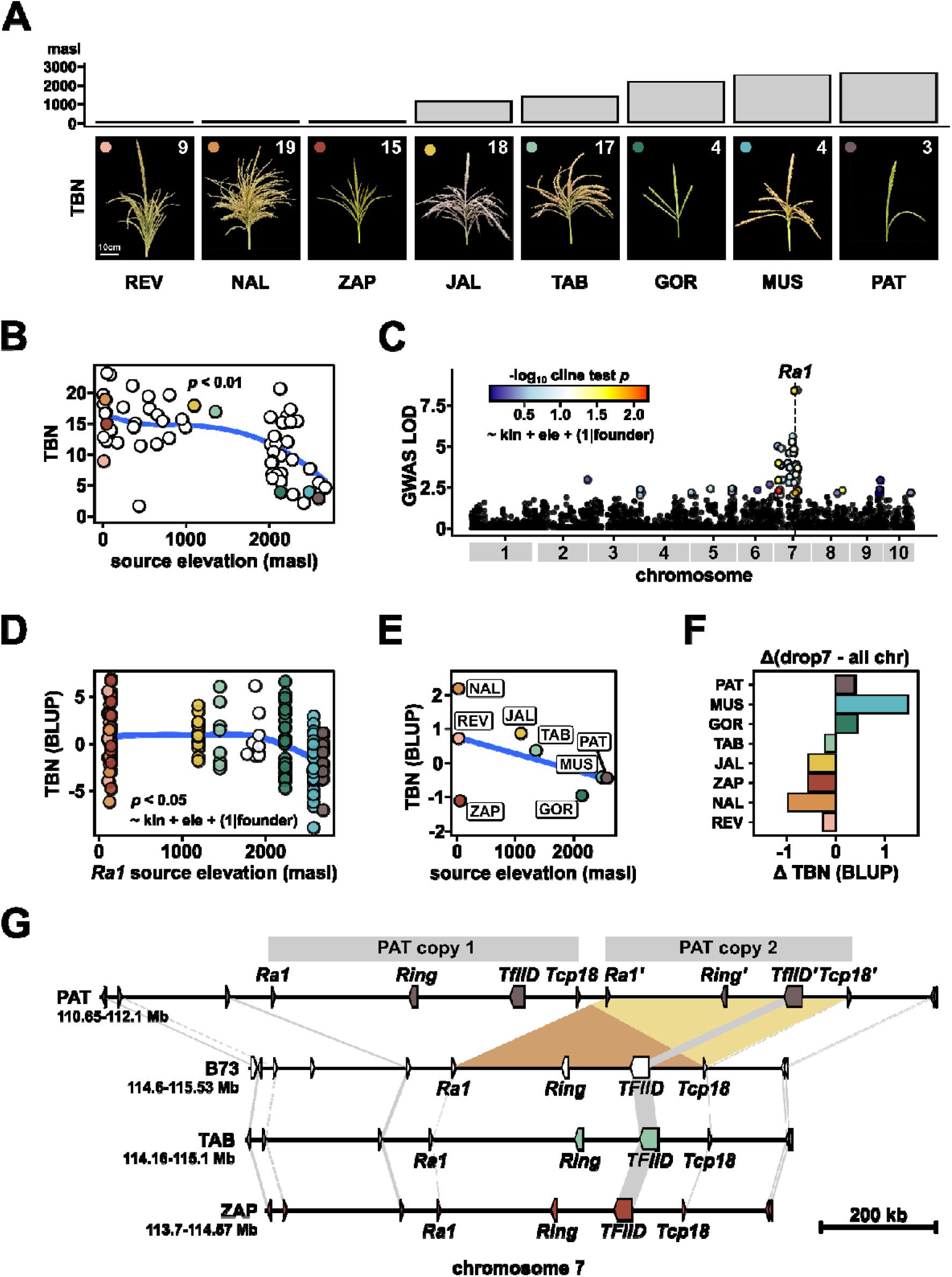
Tassel branch number follows an elevational cline in Mexican native maize. **A.** Representative tassels of the eight MexMAGIC founder accessions. The inbred founders REV, TAB and ZAP are here represented by related open-pollinated accessions. Accessions are arranged by ascending source elevation from left to right. Bars above the image show source elevation. The median tassel branch number (TBN) for each accession is shown in the upper right of the image. Colored points in the upper left of the images correspond to color legend used in other panels. **B.** Relationship between source elevation and TBN for the HiLo panel (open points) and the MexMAGIC founders (points colored as panel A). Blue line shows a LOESS fit, with the area extending +/- 1 standard error. Model statistics refer to a linear fit. **C.** GWAS support (LOD) in the MexMAGIC for association with TBN across the ten maize chromosomes. All SNPs with LOD >2 are shown. A reduced random sampling of lower-supported SNPs is shown. SNPs with LOD > 2 are colored by support for a second model (shown) fitting allele source elevation to TBN. The physical position of the candidate gene *Ra1* is indicated by a dashed vertical line. **D.** Relationship between allele source elevation at *Ra1* and TBN for the MexMAGIC families. Source elevation was calculated as the product of genotype probabilities at *Ra1* and the founder elevations. Points are colored by the founder hard-call at *Ra1* as panel A. Uncolored points represent families for which the founder call at *Ra1* was ambiguous. Blue line shows a LOESS fit, with the area extending +/-1 standard error. Model statistics refer to a linear fit following the model shown. **E.** Correlation between observed TBN and genomic BLUPs predicted by fitting marker effects from the MexMAGIC families to the founder haplotypes. Blue line shows a linear fit, with the area extending +/- 1 standard error. Statistics shown for Pearson correlation. **F.** Difference in predicted TBN for each founder between a model using all marker effects and a model dropping all chr 7 markers, including the *Ra1* QTL. **G.** Microsynteny visualization of a genome duplication in PAT around the *ra1* gene. The duplicated region spans ∼ 1.1 MB (chr 7: 114.6 - 115.53 MB in B73 ref v5) and encompasses 4 gene models including *Ra1*.

### Allele effects at a major QTL align with the elevational cline in tassel branch number

To characterize the genetic architecture of TBN in the MexMAGIC population, we evaluated 160 testcross families over two years in a mid-elevation site in western Mexico. We observed TBN values from 0 to 31 across the two years, exceeding the range seen among the eight founder varieties and the HiLo panel (Table S7). GWAS and an 8-allele QTL model identified a major QTL for TBN on chromosome (chr) 7 (Table S8), co-localized with the previously characterized TBN-associated gene *Ramosa1* (*Ra1*: Fig. 3C; Sigmon & Vollbrecht, 2010; Gonzalez-Segovia *et al*., 2019; Perez-Limón *et al*., 2022; Strable *et al*., 2023). To test alignment between allele effects and the elevational cline, we applied a second model fitting allele source elevation to TBN, taking residual population structure into account and treating founder as a random effect. We found evidence (p < 0.05) that TBN effects at *Ra1* supported the cline, driven largely by the negative effects of the MUS and PAT alleles that were sourced from the central Mexican highlands (Fig. 3C, D, S16, S17; Table S11). Interestingly, the GOR *Ra1* allele, from the highlands of northern Mexico, did not share the negative effects of MUS and PAL (Fig. 3D), although the GOR founder accession itself showed low TBN (Fig. 3A).

### A polygenic model recovers the elevational cline in TBN across founders

Although the genetic architecture of TBN was dominated by the QTL at *Ra1*, weaker signals were also found on other chromosomes (Fig. 3C; Table S9). Outside of the *Ra1* locus, top SNPs (LOD > 2) associated with a second peak on chr 7 (7_35152019) and another on chr 8 (8_160433628) showed evidence of clinal allele effects, although others, including two (6_165825744 and 8_133594448) that co-localized with previously reported TBN QTL (Xu *et al*., 2017), did not follow the elevational cline (Fig. 3C; S18; Table S12). In the latter case, it was the mid-elevation alleles that differed from both lowland and highland extremes.

Variation on a cline may be driven by selection, drift or demographic history and, in itself, does not necessarily indicate local adaptation (Savolainen *et al*., 2013; Laroche & Lenormand, 2023). Building on GEA theory and the premise of the QTL sign test (Orr, 1998), we hypothesized that a shift from predominantly positive to predominantly negative genome-wide allele effects across the cline would be consistent with directional selection on TBN. After dropping markers on chr 7, including the *Ra1* locus, SNP heritability for TBN remained at 0.33, indicating the presence of genome-wide effects contributing to TBN variation that might be used to test the hypothesis of clinal allele distribution (Fig. S19). To assess bias in the distribution of genome-wide effects with respect to the founder source, we used a polygenic model trained with the MexMAGIC families to predict TBN for the eight founder haplotypes. We found a positive correlation between observed and predicted founder TBN, and a concomitant negative correlation between predicted founder TBN and source elevation (Fig. 3E; S20), indicating that the distribution of genome-wide effects did indeed support the elevational cline. This result, however, was not robust to removal of *Ra1* and other markers on chr 7 (Fig. 3F). The reduced correlation with elevational after dropping *Ra1* was largely the result of a shift in estimate for the highland MUS founder to higher TBN, indicating the sensitivity of our analysis because of the small number of founder alleles (Fig. 3F; S21). The predicted difference between the TBN extremes NAL and PAT was preserved in the absence of *Ra1*, lending some support to the hypotheses that loci outside the *Ra1* QTL show a bias in effect at either end of the elevation cline (Fig. 3F; S21). In addition, TBN was predicted to be low in the northern highland GOR with or without inclusion of Chr 7 markers, consistent with the overall low TBN observed for this variety but the positive effect of the GOR *Ra1* allele.

### A duplication of *Ra1* in the genome of the highland variety PAT is linked to reduced TBN

To investigate possible causal variation in highland *Ra1*, we took advantage of whole genome assembly of a PAT haplotype sampled from the same accession as the highland PAT donor in the MexMAGIC, along with assemblies for the TAB (mid-elevation) and ZAP (lowland) founders (www.maizegdb.org/HiLo_project). Microsynteny analysis of the *Ra1* locus across PAT, TAB, ZAP and the B73 reference revealed a ∼1.1Mb duplication unique to PAT that contains four annotated gene models, including *Ra1* (Fig. 3G). The resulting two paralogous copies of *Ra1* in the PAT genome are 100% identical in the coding region (Table S13). Given that loss-of-function *ra1* mutants condition increased TBN (Vollbrecht *et al*., 2005; McSteen, 2006; Sigmon & Vollbrecht, 2010), we speculate that an increase in *Ra1* dosage as result of duplication in the PAT genome would reduce tassel branching. We evaluated a B73xPAT BC_3_DH NIL, derived from the sequenced PAT haplotype and carrying PAT introgression at *Ra1*, and confirmed a reduction in TBN (B73: mean = 8.3; NIL: mean = 4; Wilcoxon test for difference in means: p < 0.0001; Fig. S22).

### Highland and lowland earliness alleles segregate in the MexMAGIC population

Following TBN, we turned our attention to flowering time. In contrast to the single major QTL we found underlying TBN, prior studies lead us to anticipate that flowering variation would be driven by many smaller effect loci, offering more scope to address hypotheses related to the genetic architecture of clines. The timing of flowering is crucial to maize local adaptation (Chardon *et al*., 2004). Well studied examples include photoperiod insensitivity in temperate regions (Salvi *et al*., 2007; Ducrocq *et al*., 2008; Bouchet *et al*., 2013; Yang *et al*., 2013; Huang *et al*., 2018) and accelerated - albeit chronologically slow - development in cool highland environments within the tropics (Mercer *et al*., 2008; Romero Navarro *et al*., 2017; Janzen *et al*., 2022). Furthermore, farmers consciously maintain early and late maturing varieties that may be cultivated sequentially to ensure year round grain availability (J. J. Sanchez G. & Goodman, 1992; CONABIO, 2011) or planted simultaneously to account for climatic uncertainty (Wellhausen *et al*., 1952; Burgos-May *et al*., 2004). Although temperate photoperiod insensitivity, highland developmental acceleration and farmer-selected precocity can be characterized together as early flowering, it is unclear to what degree they are mechanistically equivalent or if they share a common genetic basis. In prior common garden evaluations across low, mid and high elevation sites in Mexico, the average flowering time (male flowering; days to anthesis, DTA) of the MexMAGIC founder varieties ranged from 60 - 91 days, the earliest being PAT, GOR (highland group) and ZAP (early maturity group; Table S1; Ruíz Corral & Sánchez González, 1998.), leading us to the expectation that both highland and lowland early flowering segregates in the MexMAGIC population. As such, we sought to assess commonalities and convergence in selection for earliness under distinct pressures.

We mapped DTA in 160 MexMAGIC test-cross families over two years in the Mexican evaluation. The broad-sense and SNP heritabilities of DTA were 0.88 and 0.79 respectively, indicating a strong genetic component. We performed a genome scan for DTA using the 8-allele model, merging the top 1% SNPs into 11 distinct QTL (Fig. 4; Table S8). There was no consistent QTL-wide trend in allele effect (we present effects accelerating flowering - *i.e.* reducing DTA - as positive) with respect to founder source (Fig. 4B, C; S23). In contrast, we observed QTL specific patterns, the QTL clustering broadly into groups driven by highland, mid-elevation and lowland earliness alleles (Fig. 4D). Applying the cline test used above for TBN, we found support (p < 0.1) for positive (*i.e.* greater flowering acceleration with increasing elevation) elevational clines at qDTA 8.1 and qDTA 8.2, and negative elevational clines at qDTA 2.3 and qDTA 9.3, indicative of distinct genetic architectures for highland and lowland precocity (Fig. 4D; S24).

**Figure 4.**
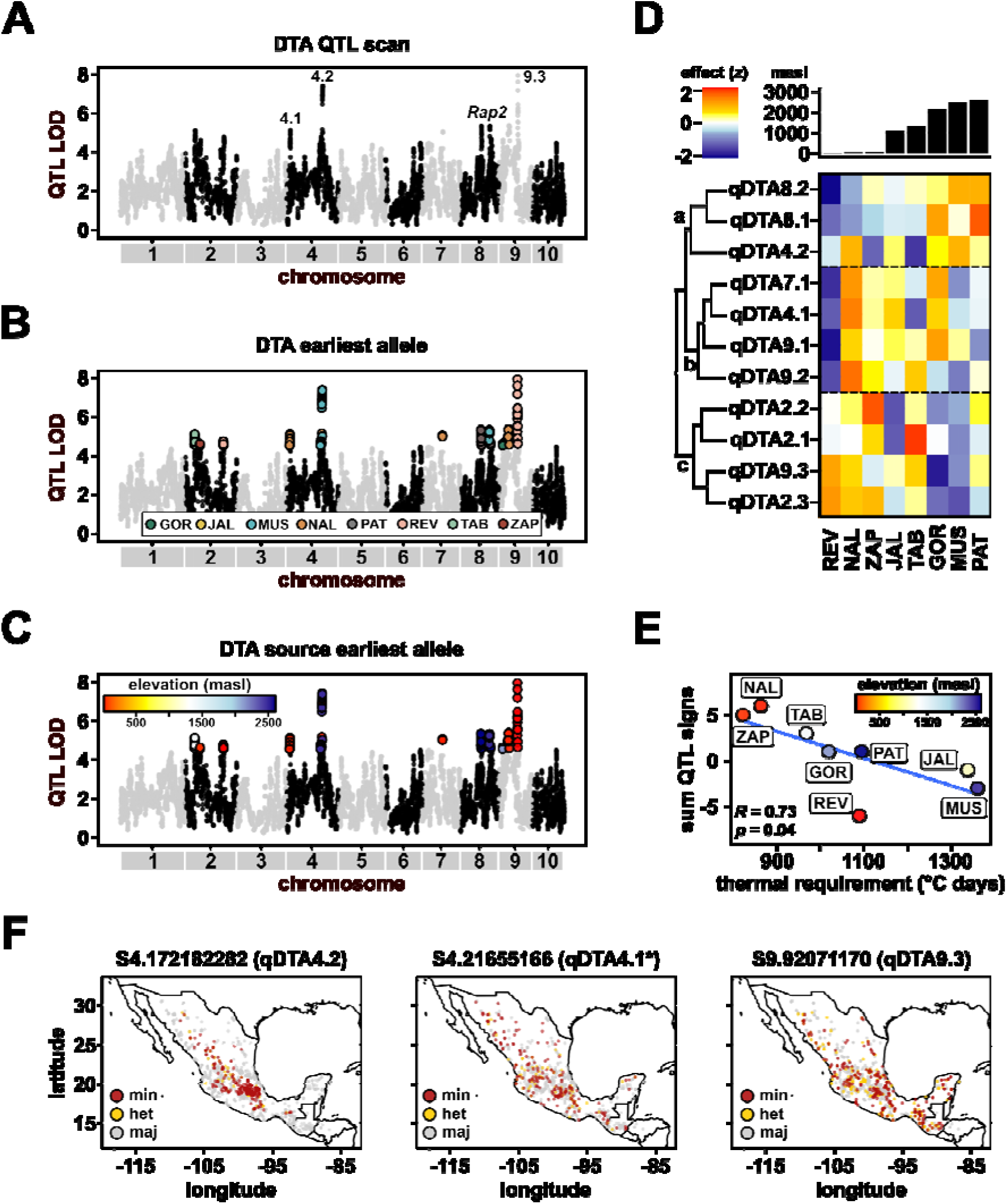
Highland and lowland earliness alleles segregate in the MexMAGIC population. **A.** SNP-level support (LOD) in the MexMAGIC for association with flowering time (days-to-anthesis, DTA) across the ten maize chromosomes. All SNPs with LOD > 4 are shown along with a reduced random sampling of lower-supported SNPs. **B, C.** As A, with top (LOD > 4) SNPs colored to indicate the founder donating the earliest allele and the source elevation of the earliest allele, respectively. **D.** Standardized allele effects (z) for each founder across DTA QTLs. Positive allele effects indicate an acceleration of flowering - *i.e.* a reduction in DTA. Founders are arranged by elevation (bar plot). QTLs are clustered by allele effect forming 3 major clusters labelled a-c for discussion in the text. **E.** Negative correlation between QTL effect sign sum and founder thermal requirement estimated by (Ruíz Corral & Sánchez González, 1998.). The color of the dot represents the founder allele source elevation **F.** Genotype in diverse Mexican maize of environmentally associated SNPs located within named MexMAGIC flowering QTL. The three QTL shown are illustrative of the three clusters shown in D. * indicates that the SNP corresponding to qDTA4.1 is located adjacent, and not within, the QTL interval.

### Earliness alleles assort preferentially into founders with a lower thermal time requirement for flowering

To investigate patterns in the accumulation of flowering alleles at the founder haplotype level, we categorized allele effects as positive (accelerating flowering), negative, or neutral, and subsequently summed the signs across the 11 QTL for each founder. A negative correlation (r^2^ = 0.53, p < 0.05) was observed between previous estimates of thermal time flowering requirement for each founder variety (Ruíz Corral & Sánchez González, 1998) and the summed allele signs across the founder haplotypes (Fig. 4E). There was no significant relationship between either thermal time requirement or QTL sign sum with elevation, consistent with the presence of early and late-flowering varieties within each elevational group and highlighting the importance of farmer selection. Indeed, although selection for accelerated flowering is a key component of highland adaptation (Eagles & Lothrop, 1994; Ruiz Corral *et al*., 2008), the lowest reported thermal time requirements and greatest accumulation of earliness alleles among the MexMAGIC founders were found in the lowland varieties NAL and ZAP.

DTA QTL intervals contained a number of compelling candidate genes, including those with homology to flowering time regulatory components in other species, candidates identified in previous analyses of selection , and hits from environmental GWA mapping (Salvi *et al*., 2007; Guo *et al*., 2018; Table S14). Nonetheless, the resolution of our mapping and limited functional characterization of potential candidates precluded confident identification of the genes underlying the QTL. That being said, we did observe overlap between qDTA8.2 and the well characterized flowering locus *Vegetative to generative transition 1* (*Vgt1*) (Salvi *et al*., 2007; Peiffer *et al*., 2014; Perez-Limón *et al*., 2022), and between qDTA4.2 and the chromosomal inversion *Inv4m* previously associated with early flowering in Mexican highland maize (Romero Navarro *et al*., 2017). To investigate evidence of our DTA variants acting in adaptation across broader Mexican maize, we identified colocalization of SNP candidates from environmental GWAS (Li *et al*., 2025) with our QTL intervals (Fig. 4E). Although no more than preliminary, it was intriguing to observe patterns of genetic stratification at environmentally associated SNPs that loosely mirrored the central highland or southern lowland origin of colocalized earliness alleles (Fig. 4E). Clearly, such an analysis would be best revisited in the future with full knowledge of the genes and variants underlying the DTA QTL.

## DISCUSSION

To address hypotheses related to the genetic architecture of locally adaptive phenotypes, we mapped two putatively adaptive traits (TBN, DTA) of differing genetic architecture in the MexMAGIC, a multiparent population designed to capture variation from across the range of Mexican native maize. The genetic architecture of TBN was dominated by a single large effect locus. In contrast, we mapped 11 QTL for DTA with evidence of distinct genetic architectures between accelerated flowering in the highlands and precocity in early-maturing lowland varieties.

Cline theory predicts that the frequency of adaptive alleles will vary with respect to changes in selective pressures across the landscape (Huxley, 1938; Haldane, 1948; Fisher, 1950). GEA inverts this expectation to infer that alleles observed to change frequency over the environment are potentially adaptative (Lasky *et al*., 2023). Although the sampling in the MexMAGIC allows only limited inference regarding allele frequency in the founder varieties, it is possible to determine to what extent allele effects are concordant with their source environment and any observed phenotypic cline. Allele effects at the TBN candidate locus *Ra1* aligned well with the elevational cline in TBN, as the two alleles sourced from the highlands of central Mexico (MUS and PAT) showed strong negative effects on TBN (Fig. 3D). Theoretically, large effect alleles in a simple genetic architecture, such as the highland *Ra1*, are more likely to align to a cline than smaller effects (Le Corre & Kremer, 2012), and, consequently, to be detectable through GEA (Lasky *et al*., 2023). Nonetheless, the strong correlation between TBN and population structure observed in the HiLo panel (Fig. S15) would likely pose a difficulty to detection of *Ra1* through GEA (Ahrens *et al*., 2021; Lasky *et al*., 2023). Indeed, we did not detect any association at *Ra1* in environmental GWAS using the CIMMYT panel (Table S14). And yet, in the context of the MexMAGIC, the correlation between source environment and *Ra1* allele effect of TBN survived correction for residual population structure, demonstrating the utility of the linkage mapping approach.

*Ra1* encodes a plant-specific C2H2 zinc finger transcription factor expressed near the base of inflorescence branch meristems to impose spikelet pair or short branch identity during ear and tassel development, respectively (Vollbrecht *et al*., 2005; McSteen, 2006; Sigmon & Vollbrecht, 2010). The timing of *Ra1* expression impacts the degree of determinacy and resulting inflorescence architecture (Strable *et al*., 2023). Loss-of-function *ra1* alleles in maize result in increased tassel branching (Vollbrecht *et al*., 2005; McSteen, 2006).

Similarly, weak *Ra1* function in the maize ancestor *Z. mays* ssp. *parviglumis* is associated with highly branched tassels (Sigmon & Vollbrecht, 2010). By analogy, we hypothesize that reduced tassel branching in Mexican highland maize is driven by an increase in *Ra1* function. The duplication of *Ra1* in a Palomero Toluqueño genome assembly (a sister haplotype to the PAT MexMAGIC founder) is clearly suggestive of a potential mechanism for increased *Ra1* activity. The adaptive significance of reduced tassel branching in the Mexican highlands has not been determined, although we note that the tassels of Mexican highland maize produce large quantities of pollen despite reduced branching (Sup. Visual 1). Although effects at *Ra1* dominated our mapping results, we did find evidence of genome wide signals supporting the TBN cline, indicative of persistent directional selection. Furthermore, although the GOR founder haplotype source from highlands northern Mexico did not carry a reduced branching *Ra1* allele, the GOR variety itself does show low tassel branching, suggesting phenotypic convergence built on a distinct genetic architecture and offering further support for adaptive significance. Such speculation would be greatly strengthened by determination of *Ra1* allelic diversity and frequency within the MexMAGIC founder varieties and more broadly in Mexican highland maize.

Maize flowering time is a heritable, complex trait governed by multiple small-effect QTL segregating across the genome (Chardon *et al*., 2004; Flint-Garcia *et al*., 2005; Buckler *et al*., 2009). Well defined latitudinal (Salvi *et al*., 2007; Ducrocq *et al*., 2008; Bouchet *et al*., 2013; Yang *et al*., 2013; Huang *et al*., 2018) and elevational (Mercer *et al*., 2008; Romero Navarro *et al*., 2017; Janzen *et al*., 2022) clines have been described in maize flowering time, blending differences in both photoperiod sensitivity and thermal requirement. Such variation is most clearly observed in a common garden (Mercer *et al*., 2008; Romero Navarro *et al*., 2017; Janzen *et al*., 2022), while the actual chronological flowering time in the native environment is strongly influenced by prevailing conditions. Beyond fulfilling the necessity of completing the cycle, maize flowering is further shaped by farmer selection for distinct maturities appropriate to different management strategies (J. J. Sanchez G. & Goodman, 1992; Wellhausen *et al*., 1952; Burgos-May *et al*., 2004; CONABIO, 2011). The MexMAGIC segregates earliness alleles contributed by both highland and lowland early maturing founders. The pattern of allelic effects we observed suggests limited molecular parallelism between accelerated highland flowering and lowland precocity, although we acknowledge the limits of our sampling. That said, major earliness alleles on chromosome 8 (here and Romero Navarro *et al*., 2017; Janzen *et al*., 2022; Perez-Limón *et al*., 2022) suggest flowering in Mexican highland maize to have more in common with early flowering in North American Flint and Northern European varieties (Ducrocq *et al*., 2008) than with early maturing Mexican lowland varieties.

The QTL qDTA4.2 co-localized with a previously described chromosomal inversion *Inv4m* that has been shown to segregate at high frequency in both Mexican highland maize and the Mexican highland teosinte *Zea mays* spp. *mexicana* (Crow *et al*., 2020; Pyhäjärvi *et al*., 2013; Calfee *et al*., 2021). The highland inverted form of *Inv4m* has been associated with a three day acceleration of flowering time in native maize (Romero Navarro *et al*., 2017). Although we linked the highland MUS and PAT alleles at qDTA4.2 to early flowering in our mid-elevation common garden, we saw similarly strong effects from the mid-elevation TAB and low-elevation NAL alleles. The Palomero Toluqueño genome assembly (a sister haplotype to the PAT MexMAGIC founder) contains the inverted allele at *Inv4m*. Furthermore, introgression analysis of PAT and MUS founders is consistent with both carrying an inverted allele at *Inv4m* as the result of gene flow from *mexicana* teosinte (Hufford *et al*., 2013; Gonzalez-Segovia *et al*., 2019). In contrast, there is no evidence for *Inv4m* in NAL, while the TAB founder genome assembly definitively carries the standard form. Collectively, our observations indicate that multiple functionally early alleles at qDTA4.2 are present in Mexican native maize, linked to both inverted and standard forms of *Inv4m*. As such, the association of early flowering with *Inv4m* may be incidental - albeit with potential consequences for subsequent selection, in contrast to a previously described example in *Mimulus* in which an inversion *per se* was the proximal cause of linked flowering acceleration (Twyford & Friedman, 2015).

In contrast to the behavior of a large effect allele, such as the highland *Ra1*, more complex genetic architectures incorporating a larger number of small allele effects exert less pressure on any given locus to align with a phenotypic cline, even if the clinal effect itself is strong (Lotterhos, 2023). Alongside *Ra1*, we detected evidence of small effect QTL associated with TBN, including those that were supported by co-localization with previously reported loci (Xu *et al*., 2017). The effects associated with minor QTL for TBN, for the most part, did not align with the elevational cline, consistent with theoretical expectation. Small effect loci likely still contribute to a phenotypic cline under the condition that the balance of positive and negative alleles carried by any given individual is appropriate to its location. Whole-genome prediction of TBN for the MexMAGIC founder haplotypes lent some support to such a model with the cline being partially maintained even after removal of strong effects at *Ra1*. In outbred populations, such as native maize varieties, a complex genetic architecture not only predicts weak patterns of changing allele frequency over a cline but would also accommodate genetic variation between phenotypically similar individuals in any specific location, provided the overall allelic complement was functionally equivalent. The flowering loci identified in the MexMAGIC showed a similar behavior as the TBN minor QTL with little discernable pattern in allele effects over the source environment. The apparently distinct genetic architectures of highland and lowland earliness segregating in the MexMAGIC further complicated efforts at simple alignment of allele effects to a single environmental axis. Nonetheless, whole-genome prediction based on founder haplotypes was consistent with previously reported thermal time requirement for the founder varieties.

The traits we have characterized in the MexMAGIC are components of local adaptation. The genetic architecture of local adaptation itself, including all the components that define “fitness” for a crop species, is necessarily far more complex. Taking into account the possibility of pleiotropy and epistatic interactions, the challenge to GEA approaches is clear. Further evaluation of resources such as the MexMAGIC, notably in different environments, will provide a valuable complement to GEA and genetic differentiation methods in efforts to understand the basis of local adaptation with a view to conservation and responsible use of native biodiversity resources.

## MATERIALS AND METHODS

### Selection of MexMAGIC founders

To capture haplotypes sourced from across the environmental range of Mexican native maize, eight accessions were selected corresponding to varieties placed in different climatic-adaptation groups in a previous comprehensive analysis of 42 native varieties (Eagles & Lothrop, 1994; Ruiz Corral *et al*., 2008). Three varieties (REV, TAB, and ZAP) were represented by inbred (S_6_) derivatives of outbred accessions (Chia *et al*., 2012; Venkatesh *et al*., 2015), and the other five varieties (GOR, JAL, MUS, NAL and PAL) by outbred open pollinated varieties (OPVs) obtained from the International Center for Maize and Wheat Improvement (CIMMyT) seed bank (Table S1; CIMMYTMA-007937, CIMMYTMA-002246, CIMMYTMA-005582, CIMMYTMA-000815, CIMMYTMA-002233). The accession of the highland variety PAL was selected because of previous genomic (Vielle-Calzada *et al*., 2009) and genetic (Perez-Limón *et al*., 2022) characterization. Accessions of JAL and NAL were selected as members of a previously characterized core diversity set (Wellhausen *et al*., 1952; Reif *et al*., 2006), although subsequent classification has questioned whether the accession Nayarit 6 (CIMMYTMA-002246) is representative of the Jala variety (Denise Costich & Cristian Zavalla Espinosa, unpublished observation). The varieties GOR and MUS were selected to represent the unique environments of Chihuahua and Michoacán, respectively.

### Environmental analysis of the Mexican diversity panel

To provide context for our selected MexMAGIC founders, we defined a broader Mexican diversity panel from a collection of georeferenced and genotyped native maize varieties previously characterized by CIMMYT (Romero Navarro *et al*., 2017). After filtering for Mexican origin (latitude ranging from 14.38 to 30.64; longitude ranging from -111.12 to - 86.92; three Guatemalan accessions were retained), availability of genotype data, and unambiguous variety assignment, we obtained a set of 1,379 accessions (Table S2).

Georeference data was used to associate climate data for each accession from the WorldClim Database (Hijmans *et al*., 2005) and the Global Soil Dataset for use in Earth System Models (GSDE; Shangguan *et al*., 2014) using {R/raster::extract} (Hijmans, 2023) as previously described (Lasky *et al*., 2015; McLaughlin *et al*., 2024). Environmental descriptors selected for further analysis were annual and seasonal mean temperature and precipitation, mean temperature of the coldest month, soil pH, and seasonal average photoperiod. To calculate seasonal means, the growing season was approximated as May to October for all varieties and climatic zones. Soil pH was calculated as the mean of pH-H_2_O, pH-KCl and pH CaCl_2_ descriptors from GSDE. Photoperiod was calculated from latitude using {R/geosphere::daylength} (Hijmans, 2022). Accessions were assigned to varieties using the GrinGlobal database (https://www.grin-global.org) and subsequently matched by variety name to climatic-adaptation groups based on a previously published classification (Eagles & Lothrop, 1994; Ruiz Corral *et al*., 2008) and plotted on the map of Mexico using {R/rworldmap::map.polygon} (South, 2011). Environmental principal component (PC) analysis was performed on the MexMAGIC founders and Mexican diversity panel accessions with the eight environmental descriptors and elevation as variables, using {R/ade4::dudi.pca} (Dray & Dufour, 2007). Variables were centered and scaled for analysis.

### Genome Wide Association Using Environmental Descriptors

Environmental GWAS was performed using genotyping-by-sequencing data (GBS) from the 1,379 accessions of the Mexican HiLo panel and 8 environmental descriptors (Table S2) using a previously described pipeline (Li *et al*., 2025). Briefly, environmental descriptors were rank inverse normal transformed and GBS SNPs were filtered to a minor allele frequency > 0.01 for a total of 343,205 markers. To perform multi-trait GWAS on multiple environmental variables, the package MegaLMM (Runcie *et al*., 2021) was first run with a kinship matrix generated from the GBS data and all transformed environmental variables, and the de-correlated genetic and residual covariance matrices included as structured error terms in an EMMAX ANOVA model implemented in the package JointGWAS. To identify potential candidate genes from MEGALMM environmental GWAS analysis, the SNP-based evidence of association (p-values) were collapsed into gene-based evidence using MAGMA (de Leeuw *et al*., 2015). After an FDR correction (Storey *et al*., 2025), only the top 1% significant genes were kept (Table S14).

### Generation of MexMAGIC families

Crosses among the eight selected MexMAGIC founders (5 outbred varieties; 3 S_6_ inbred lines) were attempted following a diallel scheme and a set of four successful F_1_ combinations selected to cover all eight parents. Subsequently, four-way crosses were made using a single individual from each F_1_ to capture a single haplotype per founder using the “chain” scheme A to B, B to C, C to D and D to A. An equal number of seeds was advanced from each four-way ear for a round of eight-way crosses using multiple individuals, followed by 3 generations of random intermating, and 3 generations of inbreeding. For simplicity, the final families were designated Intermated3 Selfed3 (I_3_S_3_). The I_3_S_3_ MexMAGIC were test-crossed to the inbred line NC358 for following field evaluations. More details of population generation are given in Supplemental Material.

### Determination of MexMAGIC founder haplotypes

To define the eight founder haplotypes of the MexMAGIC, we generated Whole Genome Sequence (WGS) at ∼33x coverage for the four F_1_ individuals (Figure 1), two S_6_ founders (Tabloncillo and Reventador) and OPV accessions of Jala and Palomero Toluqueño, as well as generating full genome assemblies of Zapalote Chico and Tabloncillo (www.maizegdb.org/HiLo_project). To identify genetic variants segregating in the population, we used the GATK Best Practices workflow for Germline short variant discovery (Poplin *et al*., 2018) for both WGS and full genome assemblies. Initial quality assessment and preprocessing of WGS data was performed under default parameters using tools described in https://bioinformaticsworkbook.org/dataAnalysis/VariantCalling/gatk-dnaseq-best-practices-workflow.html#gsc.tab=0. Short reads were aligned to the B73 genome V4 (Jiao *et al*., 2017) using bwa (Li & Durbin, 2009). A first round of variant discovery from BAM files was performed using {gatk::HaplotypeCaller}. Identified variants were filtered by quality values, and used as a training set to recalibrate the BAM files using {gatk::BaseRecalibrator}. A second round of variant discovery was performed in the re-calibrated BAM files using {gatk::HaplotypeCaller}, resulting in a GVCF. The full genome assemblies of Zapalote Chico and Tabloncillo were aligned to the B73 v4 reference genome using the {AnchorWave::genoAli} function (Song *et al*., 2022) with -IV parameter to detect inversions and other options as default as suggested in https://github.com/baoxingsong/AnchorWave. The resulting MAF files were converted to GVCF using the {TASSEL5::MAFToGVCFPlugin} plugin (Bradbury *et al*., 2007) with default parameters according to https://github.com/baoxingsong/AnchorWave/blob/master/doc/GATK.md. The GVCFs generated for the WGS samples and full genome assemblies were combined into a single database using {gatk::GenomicsDBImport}, and a joint calling on the full cohort was performed with {gatk::GenotypeGVCFs}. The resulting VCFs were hard filtered using default parameters as described in https://gatk.broadinstitute.org/hc/en-us/articles/360035890471-Hard-filtering-germline-short-variant. SNPs with missing information were removed, resulting in a final VCF containing ∼ 16 million SNPs distributed across the 10 maize chromosomes. VCFs were transformed into HAPMAP format using TASSEL5 (Bradbury *et al*., 2007) and passed to a custom script in {R} (R Core Team, 2022) that implemented a triplet approach comparing parents and their F1 to estimate the allele contributed by each of the two parents at sites heterozygous in the F1 cross. For the crosses TAB x MUS, GOR x REV and NAL x ZAP, one of the two parents was represented by an S_6_ partial inbred. For the Cross JAL x PAT, both parents were OPVs (Fig. 2A). Sites that could not be resolved were dropped, yielding a final set of ∼ 8 million pseudo-homozygous SNPs for the eight founders of the MexMAGIC population.

### PacBio HiFi LRS sequencing

PacBio HiFi-grade genomic DNA (gDNA) was isolated from approximately 0.5 g of frozen young leaf tissue from single plants using the NucleoBond® HMW DNA kit (Cat. 740160.20; Macherey-Nagel; Allentown, PA, USA) following the manufacturer’s instructions. Native gDNA was sheared to a 10–20 kb fragment size distribution using a Diagenode Megaruptor 3, and fragment size was assessed with the Agilent Femto Pulse System. Fragments shorter than 10 kb were removed using the PippinHT system (Sage Science; Beverly, MA, USA). Sequencing libraries were prepared with the SMRTbell Express Template Prep Kit 2.0 (Pacific Biosciences; California, USA) and sequenced using Chemistry 2.0 on the PacBio Sequel II platform, following standard protocols. Data was collected from two SMRT Cells per sample in 30-hour movies. HiFi reads shorter than 5 kb were filtered in silico after sequencing. For the Tabloncillo accession, 4,438,228 reads totaling 72.93 Gb (∼34× coverage) with an average read length of 16.4 kb were obtained. For Zapalote Chico, 4,745,054 reads totaling 71 Gb with an average read length of 15.9 kb were collected, and for Palomero de Jalisco, 5,529,410 reads totaling 63.7 Gb with an average read length of 11.8 kb were obtained. For Palomero Toluqueño 38x with a read N50 of 14.5 kb was obtained.

### Bionano DLS Optical genome mapping

Ultra-high molecular weight (uHMW) DNA extraction and optical genome mapping using the Bionano Direct Label and Stain (DLS) chemistry were performed as described in Hufford et al (2021). The resulting molecule datasets were filtered to retain only molecules larger than a minimum length threshold needed to reach a total raw molecule coverage of approximately 150-180x. For Tabloncillo, 998,133 molecules ≥350 kb were collected, totaling 491 Gb. For Zapalote Chico, 1,179,310 molecules ≥300 kb were obtained, totaling 498 Gb. For Palomero de Jalisco, 1,606,337 molecules ≥150 kb were collected, totaling 409 Gb. Bionano Access v.1.7.2 and Bionano Tools v3.7 were used to process, visualize, assemble and scaffold maps. Zapalote Chico assembly produced 86 maps with a N50 length of 89 Mb and total length of 2,147 Mb. Palomero de Jalisco assembled into 87 maps with an N50 length of 109.7 Mb and total length of 2,428.78. The assembly of Tabloncillo led to 69 maps with N50 length of 99.74 and 2,217 Mbp of total length.

### Genotypic analysis of MexMAGIC families

Ten seeds each of 221 MexMAGIC families I_3_S_3_ families and the MexMAGIC founders were germinated in the greenhouse and tissue collected for genotypic analysis. For MexMAGIC families, tissue of 10 individuals was pooled. For MexMAGIC founders, individuals were analyzed separately. Leaf tissue was pulverized to a fine powder using a mortar and pestle in liquid N_2_. DNA was extracted from frozen tissue using the DNeasy Plant Kit (Qiagen). A total of 240 samples were analyzed using the Illumina MaizeLD BeadChip (https://www.illumina.com/documents/products/datasheets/datasheet_maize_snp50.pdf) by the Genomic Services Laboratory at the Advanced Genomics Unit, Irapuato, Mexico. Polymorphic sites with a call rate >95% were used for further analysis, yielding approximately 20,000 markers.

### Generation of a MexMAGIC Practical Haplotype Graph

To impute founder haplotypes onto the chip genotyped MexMAGIC families, we built a custom Practical Haplotype Graph (PHG, (Bradbury *et al*., 2022). The PHG was built with 1000 intervals (100 per chromosome) and populated using the 8 founder haplotypes and the B73 reference (Jiao *et al*., 2017). The founder haplotype HAPMAP file was transformed to VCF using TASSEL5 and then converted to BAM using {gatk::SimulateReadsForVariants} and the finally to GVCF using {gatk::HaplotypeCaller}. The resulting GVCFs were used to populate the PHG and to impute onto the MexMAGIC families genotyped with the Illumina MaizeLD BeadChip. Imputation increased the number of SNPs per family from ∼ 10,000 to ∼ 6M SNPs. The imputed set was filtered by minor allele frequency consistent with the expected frequency across the founders, generating a dataset of ∼3.5 M SNPs. To estimate the relative contribution of each founder to the population, the founder assigned to each range in each family was extracted using the {R/rPHG} package (Monier *et al*., 2024). The genome-wide founder and missing data proportion was estimated as the count of ranges assigned to the founder across the complete population (200 families) divided by the total number of ranges (1000 ranges per family).

### Population structure of MexMAGIC families

The filtered set of 10,191 BeadChip SNPs for MexMAGIC families and founder parents was converted to numeric format using TASSEL and missing marker data was imputed using {R/rrBLUP::A.mat} (Endelman, 2011) under default parameters. Population structure was estimated as the first five principal coordinates (PCoAs) obtained from multidimensional scaling of the genotype matrix using {R/stats::cmdscale}. The relationship between PCo2 and source elevation for the founder parents was modeled using {R/stats::loess} with a span of 0.4 determined by inspection, and theoretical elevations for the MexMAGIC families obtained from the loess fit using {R/stats::predict}.

### Parental assignment across the genomes of MexMAGIC families

To characterize the genetic mosaic of the MexMAGIC families, a subset of 1M random SNPs was selected with stratification by chromosome using the {R/rsample::initial_split} function (Frick *et al*., 2024). To give a genetic position to SNPs in the MexMAGIC dataset, we trained a monotonic smooth using {R/scam} package (Pya & Wood, 2015) to estimate the relationship between the physical and genetic position of the SNPs in the IBM RIL maize population (Lee *et al*., 2002), and used it to predict the genetic position on the MexMAGIC dataset. The genetic and physical map, and founder and MexMAGIC families genotypes were combined into a cross object to be used by the {R/qtl2} package. SNPs with identical genotype data were identified with the {qtl2::find_dup_markers} function and dropped, generating a final dataset of 17,030 markers. The founder conditional genotype probabilities at every marker were estimated with the {qtl2::calc_genoprob} function and the founder genotype with maximum marginal probability was estimated with the {qtl2::maxmarg} function (Figure 2E).

### Field evaluation of MexMAGIC families

MexMAGIC families were evaluated as test-crosses to the inbred line NC350 in the summers of 2021 and 2022 in Ameca, Jalisco, Mexico (20.575024, - 104.038857; 1235 m.a.s.l.). Two complete blocks were evaluated in 2021 and 2022, consisting of 190 and 175 families, respectively. The distribution of the families was completely randomized within each block. 15 seeds per family were sown in 2 m long rows. Plants were grown under conventional agronomic conditions. A commercial hybrid adapted to the region was planted as a check across the field (Fig. S11, S12).

### Evaluation of an elevational cline in tassel branch number

Tassel branch number (TBN), source elevation and genotype data for 52 Mexican native maize accessions (HiLo panel) was obtained from (Janzen *et al*., 2022). The published sampling was stratified by elevation, with paired “high” and “low” accessions selected along a latitudinal gradient from northern to southern Mexico. TBN data were summarized as the median of values reported for highland and lowland common gardens (Sup. Table 10). The published genotypic data was filtered by minor allele frequency > 0.1 and missing information < 0.2, to 4060 SNPs using TASSEL5 (Bradbury *et al*., 2007) and converted to numeric format (Table S15). Missing marker data was imputed using {R/rrBLUP::A.mat} (Endelman, 2011) under default parameters. Population structure for the Janzen et al. (2022) data was estimated as the first five principal coordinates (PCoAs) obtained from multidimensional scaling of the genotype matrix using {R/stats::cmdscale} (Fig. S14). The relationship between source elevation and TBN was investigated using {R/stats::lm} to fit the fixed effect models:

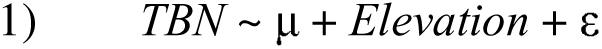

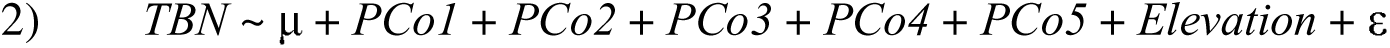

Although highly significant in the first model (Eq. 1; *p* < 0.001), the elevation effect on TBN did not survive inclusion of population PCo1-5 before elevation in the second model (Eq. 2; Type II ANOVAelevation *p* = 0.99), reflecting the structuring of the panel by elevation and the correlation between PCo1 and elevation (Fig. S15).

### Mapping in the MexMAGIC population

To map functional variation in the MexMAGIC founders, we used two different mapping methodologies in the {R/qtl2} package (Broman *et al*., 2019): SNP-association and 8-allele QTL mapping. Both methods use a mixed linear model to model the phenotypic variation as a function of the genetic variation in the population, with the addition of a kinship matrix to account for residual population structure. The SNP-Association method, collapsed the marginal founder probabilities into biallelic allele probabilities and performed the association at two different genetic levels; the 8-allele QTL model, in contrast, directly used the marginal founder conditional probabilities, performing the association at eight different levels corresponding to the eight founders. A “leave one chromosome out” (LOCO) kinship matrix was estimated using {qtl2::calc_kinship()} function, and the previously founder conditional genotype probabilities were turned into allele probabilities using {qtl2::genoprob_to_alleleprob()} function. A linear mixed model was fitted using the {qtl2::scan1snps()} function, using as inputs the allele probabilities, phenotypic information, and the kinship matrix to account for residual population structure. The results of the SNP-association were summarized in Manhattan plots using {R/ggplot2} package for R (Wickham, 2016). To understand the effect of the association between the segregation of phenotypes and the effect of the founder alleles, a single-QTL scan was performed using an 8-allele method. A linear mixed model was fitted using the {qtl2::scan1} function, using the marginal genotype probabilities, phenotypic information and the kinship matrix. The LOD trace plots were visualized using the {R/ggplot2} package. To establish a threshold to detect significant signals, a 1000 permutation test was run using the function {qtl2::scan1perm} function considering a significance threshold (alpha) of 0.1. Significant QTLs were identified with the function {qtl2::find_peaks} and the LOD-support interval was considered as a drop of 2 in LOD value from the QTL peak (Broman *et al*., 2003). To obtain an estimation and confidence interval of the allelic founder effect at every QTL, the marginal genotype probabilities at the lead SNP were extracted using {ql2::pull_genoprobpos}. The subset of genotype probabilities, kinship matrix and phenotypic data were used as input for the {qtl2::fit1} function to fit a single-QTL model using a mixed linear model and kinship matrix to control for residual structure. The estimated coefficient and standard errors of the founder allelic effect were extracted from the QTL object for analysis and visualization.

### Test of environmental clines in the MexMAGIC population

To test for alignment between allele effects at candidate SNPs and elevational, a mixed linear model was fitted using {R/lme4} and {R/lmerTest}(Bates *et al*., 2007; Kuznetsova *et al*., 2017).

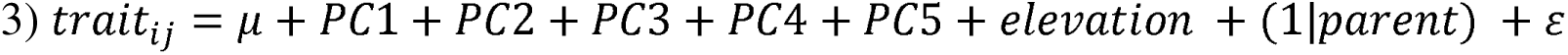

We modeled trait variation as a function of the fixed effects of the first five genetic PCos, source elevation, and the random effect of the hard call on founder genotype (Equation 3). The marginal genotype probabilities at a candidate SNP was extracted using {ql2::pull_genoprobpos} function, and the hard call for every family (genotype with the maximum marginal probability with a confidence of at least 0.9) was estimated with the {qtl2::maxmarg}; the hard call for a family with a genotype confidence smaller than 0.9 was considered as missing . To make use of observations without a hard call, the elevation was estimated as the dot product between the marginal genotype probabilities and the founder-allele source elevation. The significance of the model was extracted from the resulting object of the {lmerTest::anova()}function.

## GENOME ACCESSION NUMBERS

PAT: ERS16235157, GCA_963555555; TAB: ERS16997546, GCA_964199785; ZAP: ERS16997547; GCA_964199775

## ACKNOWLEDGEMENTS

We are indebted to many members of the former Maize Genetics and Genomics Group at the Unidad de Genómica Avanzada, Irapuato, who contributed to the generation and evaluation of the material described in this work, including Jorge-Vladimir Torres-Rodriguez, Fausto Rodríguez-Zapata, Benjamín Barrales-Gamez, Dario Alavez-Mercado and Fabian-Oswaldo Santa Maria-Gasca. We thank Cruz Robledo and his team (Puerto Vallarta Agricultural Invernal), and Beda Angehrn and Mario Rivera (UNISEM) for their invaluable assistance in genetic nurseries and field evaluation. This work would not have been possible without the international maize research community and the willingness of so many colleagues to support the development of the Irapuato research program. We thank Sherry Flint-Garcia, Matthew Hufford, Denise Costich and Cristian Zavalla Espinosa for support and valuable discussions. Finally, we would like to acknowledge the smallholder farmers and indigenous people of the Americas whose work and love for their traditions and identity keep maize diversity alive.

## AUTHOR CONTRIBUTIONS

RJHS, RRA and CSG designed the research; MNSV, ALAN, MRRF, CCM, JCM, SPL, CSG, RRA and RJHS generated and evaluated the MexMAGIC population; MGP, OHSP and JSS collected and analyzed additional field data; MRRF and PY performed genotypic analysis; VL and KAF performed genome assembly; FL, DER and JRI performed environmental association analysis; SPL, RJHS, PY, DER and JRI conducted primary analysis and interpretation of results; all authors contributed to writing the manuscript.

## REFERENCES

1. Ahrens CW, Jordan R, Bragg J, Harrison PA, Hopley T, Bothwell H, Murray K, Steane DA, Whale JW, Byrne M, et al. 2021. Regarding the F-word: The effects of data filtering on inferred genotype-environment associations. Molecular ecology resources 21: 1460–1474.

2. Bates D, Sarkar D, Bates MD, Matrix L. 2007. The lme4 package. R package version 2: 74.

3. Bouchet S, Servin B, Bertin P, Madur D, Combes V, Dumas F, Brunel D, Laborde J, Charcosset A, Nicolas S. 2013. Adaptation of maize to temperate climates: mid-density genome-wide association genetics and diversity patterns reveal key genomic regions, with a major contribution of the Vgt2 (ZCN8) locus. PloS one 8: e71377.

4. Bradbury PJ, Casstevens T, Jensen SE, Johnson LC, Miller ZR, Monier B, Romay MC, Song B, Buckler ES. 2022. The Practical Haplotype Graph, a platform for storing and using pangenomes for imputation. Bioinformatics 38: 3698–3702.

5. Bradbury PJ, Zhang Z, Kroon DE, Casstevens TM, Ramdoss Y, Buckler ES. 2007. TASSEL: software for association mapping of complex traits in diverse samples. Bioinformatics 23: 2633–2635.

6. Broman KW, Gatti DM, Simecek P, Furlotte NA, Prins P, Sen Ś, Yandell BS, Churchill GA. 2019. R/qtl2: Software for mapping quantitative trait loci with high-dimensional data and multiparent populations. Genetics 211: 495–502.

7. Broman KW, Wu H, Sen S, Churchill GA. 2003. R/qtl: QTL mapping in experimental crosses. Bioinformatics 19: 889–890.

8. Buckler ES, Holland JB, Bradbury PJ, Acharya CB, Brown PJ, Browne C, Ersoz E, Flint-Garcia S, Garcia A, Glaubitz JC, et al. 2009. The genetic architecture of maize flowering time. Science 325: 714–718.

9. Buckner B, Miguel PS, Janick-Buckner D, Bennetzen JL. 1996. The y1 gene of maize codes for phytoene synthase. Genetics 143: 479–488.

10. Burgos-May LA, Chávez-Servia JL, Ortiz-Cereceres J. 2004. Variabilidad morfológica de maíces criollos de la península de Yucatán, México. In: Chávez-Servia JL, Tuxill J, Jarvis DI, eds. Manejo de la diversidad de los cultivos en los agroecosistemas tradicionales. Cali, Colombia: Instituto Internacional de Recursos Fitogenéticos, 58–66.

11. Calfee E, Gates D, Lorant A, Perkins MT, Coop G, Ross-Ibarra J. 2021. Selective sorting of ancestral introgression in maize and teosinte along an elevational cline. PLoS genetics 17: e1009810.

12. Campbell Q, Bedford JA, Yu Y, Halpin-McCormick A, Castaneda-Alvarez N, Runck B, Neyhart J, Ewing P, Ortiz-Barrientos D, Gao L, et al. 2025. Agricultural landscape genomics to increase crop resilience. Plant communications 6: 101260.

13. Chardon F, Virlon B, Moreau L, Falque M, Joets J, Decousset L, Murigneux A, Charcosset A. 2004. Genetic architecture of flowering time in maize as inferred from quantitative trait loci meta-analysis and synteny conservation with the rice genome. Genetics 168: 2169–2185.

14. Chen W, Cui F, Zhu H, Zhang X, Lu S, Lu C, Chang H, Fan L, Lin H, Fang J, et al. 2024. Genome-wide association study of kernel colour traits and mining of elite alleles from the major loci in maize. BMC plant biology 24: 25.

15. Chia J-M, Song C, Bradbury PJ, Costich D, de Leon N, Doebley J, Elshire RJ, Gaut B, Geller L, Glaubitz JC, et al. 2012. Maize HapMap2 identifies extant variation from a genome in flux. Nature genetics 44: 803–807.

16. Cleveland DA, Soleri D. 2007. Extending Darwin’s Analogy: Bridging Differences in Concepts of Selection between Farmers, Biologists, and Plant Breeders. Economic botany 61: 121–136.

17. CONABIO. 2011. Base de datos del proyecto global ‘Recopilación, generación, actualización y análisis de información acerca de la diversidad genética de maíces y sus parientes silvestres en México’.

18. Crow T, Ta J, Nojoomi S, Aguilar-Rangel MR, Torres Rodríguez JV, Gates D, Rellán-Álvarez R, Sawers R, Runcie D. 2020. Gene regulatory effects of a large chromosomal inversion in highland maize. PLoS genetics 16: e1009213.

19. Dray S, Dufour A-B. 2007. Theade4Package: Implementing the duality diagram for ecologists. Journal of statistical software 22.

20. Ducrocq S, Madur D, Veyrieras J-B, Camus-Kulandaivelu L, Kloiber-Maitz M, Presterl T, Ouzunova M, Manicacci D, Charcosset A. 2008. Key impact of Vgt1 on flowering time adaptation in maize: evidence from association mapping and ecogeographical information. Genetics 178: 2433–2437.

21. Eagles HA, Lothrop JE. 1994. Highland Maize from Central Mexico—Its Origin, Characteristics, and Use in Breeding Programs. Crop science 34: 11–19.

22. Endelman JB. 2011. Ridge regression and other kernels for genomic selection with R package rrBLUP. The plant genome 4: 250–255.

23. Fisher RA. 1950. Gene frequencies in a cline determined by selection and diffusion. Biometrics 6: 353–361.

24. Flint-Garcia SA, Thuillet A-C, Yu J, Pressoir G, Romero SM, Mitchell SE, Doebley J, Kresovich S, Goodman MM, Buckler ES. 2005. Maize association population: a high-resolution platform for quantitative trait locus dissection: High-resolution maize association population. The Plant journal: for cell and molecular biology 44: 1054–1064.

25. Frick H, Chow F, Kuhn M, Mahoney M, Silge J, Wickham H. 2024. rsample: General Resampling Infrastructure.

26. Gonzalez-Segovia E, Pérez-Limon S, Cíntora-Martínez GC, Guerrero-Zavala A, Janzen GM, Hufford MB, Ross-Ibarra J, Sawers RJH. 2019. Characterization of introgression from the teosinte ssp. to Mexican highland maize. PeerJ 7: e6815.

27. Grotewold E, Drummond BJ, Bowen B, Peterson T. 1994. The myb-homologous P gene controls phlobaphene pigmentation in maize floral organs by directly activating a flavonoid biosynthetic gene subset. Cell 76: 543–553.

28. Guo L, Wang X, Zhao M, Chen Q, Doebley JF, Tian F, Huang C, Li C, Li D, Yang CJ, et al. 2018. Stepwise cis-Regulatory Changes in ZCN8 Contribute to Maize Flowering-Time Adaptation | Elsevier Enhanced Reader. Current biology: CB: 3005–3015.

29. Haldane JBS. 1948. The theory of a cline. Journal of genetics 48: 277–284.

30. Hijmans RJ. 2022. geosphere: Spherical Trigonometry.

31. Hijmans RJ. 2023. raster: Geographic Data Analysis and Modeling.

32. Hijmans RJ, Cameron SE, Parra JL, Jones PG, Jarvis A. 2005. Very high resolution interpolated climate surfaces for global land areas. International Journal of Climatology 25: 1965–1978.

33. Huang C, Sun H, Xu D, Chen Q, Liang Y, Wang X, Xu G, Tian J, Wang C, Li D, et al. 2018. ZmCCT9 enhances maize adaptation to higher latitudes. Proceedings of the National Academy of Sciences of the United States of America 115: E334–E341.

34. Hufford MB, Lubinksy P, Pyhäjärvi T, Devengenzo MT, Ellstrand NC, Ross-Ibarra J. 2013. The genomic signature of crop-wild introgression in maize. PLoS genetics 9: e1003477.

35. Huxley J. 1938. Clines: An auxiliary taxonomic principle. Nature 142: 219–220.

36. Janzen GM, Aguilar-Rangel MR, Cíntora-Martínez C, Blöcher-Juárez KA, González-Segovia E, Studer AJ, Runcie DE, Flint-Garcia SA, Rellán-Álvarez R, Sawers RJH, et al. 2022. Demonstration of local adaptation in maize landraces by reciprocal transplantation. Evolutionary applications 15: 817–837.

37. Jiao Y, Peluso P, Shi J, Liang T, Stitzer MC, Wang B, Campbell MS, Stein JC, Wei X, Chin C-S, et al. 2017. Improved maize reference genome with single-molecule technologies. Nature 546: 524–527.

38. J. J. Sanchez G., Goodman MM. 1992. Relationships among the Mexican races of maize. Economic botany 46: 72–85.

39. Key FM, Abdul-Aziz MA, Mundry R, Peter BM, Sekar A, D’Amato M, Dennis MY, Schmidt JM, Andrés AM. 2018. Human local adaptation of the TRPM8 cold receptor along a latitudinal cline. PLoS genetics 14: e1007298.

40. Koski MH, Ashman T-L. 2015. An altitudinal cline in UV floral pattern corresponds with a behavioral change of a generalist pollinator assemblage. Ecology 96: 3343–3353.

41. Kuznetsova A, Brockhoff PB, Christensen RHB. 2017. LmerTest package: Tests in linear mixed effects models. Journal of statistical software 82: 1–26.

42. Laroche F, Lenormand T. 2023. The genetic architecture of local adaptation in a cline. Peer community journal 3.

43. Lasky JR, Josephs EB, Morris GP. 2023. Genotype-environment associations to reveal the molecular basis of environmental adaptation. The Plant cell 35: 125–138.

44. Lasky JR, Upadhyaya HD, Ramu P, Deshpande S, Hash CT, Bonnette J, Juenger TE, Hyma K, Acharya C, Mitchell SE, et al. 2015. Genome-environment associations in sorghum landraces predict adaptive traits. Science advances 1: e1400218.

45. Le Corre V, Kremer A. 2012. The genetic differentiation at quantitative trait loci under local adaptation. Molecular ecology 21: 1548–1566.

46. Lee M, Sharopova N, Beavis WD, Grant D, Katt M, Blair D, Hallauer A. 2002. Expanding the genetic map of maize with the intermated B73 x Mo17 (IBM) population. Plant molecular biology 48: 453–461.

47. de Leeuw CA, Mooij JM, Heskes T, Posthuma D. 2015. MAGMA: generalized gene-set analysis of GWAS data. PLoS computational biology 11: e1004219.

48. Li H, Durbin R. 2009. Fast and accurate short read alignment with Burrows-Wheeler transform. Bioinformatics (Oxford, England) 25: 1754–1760.

49. Li F, Gates DJ, Buckler ES, Hufford MB, Janzen GM, Rellán-Álvarez R, Rodríguez-Zapata F, Romero Navarro JA, Sawers RJH, Snodgrass SJ, et al. 2025. Environmental data provide marginal benefit for predicting climate adaptation. PLoS genetics 21: e1011714.

50. Lotterhos KE. 2023. The paradox of adaptive trait clines with nonclinal patterns in the underlying genes. Proceedings of the National Academy of Sciences of the United States of America 120: e2220313120.

51. Louette D, Charrier A, Berthaud J. 1997. In Situ conservation of maize in Mexico: Genetic diversity and Maize seed management in a traditional community. Economic botany 51: 20–38.

52. Mao D, Xin Y, Tan Y, Hu X, Bai J, Liu Z-Y, Yu Y, Li L, Peng C, Fan T, et al. 2019. Natural variation in the HAN1 gene confers chilling tolerance in rice and allowed adaptation to a temperate climate. Proceedings of the National Academy of Sciences of the United States of America 116: 3494–3501.

53. Matsuoka Y, Vigouroux Y, Goodman MM, Sanchez G J, Buckler E, Doebley J. 2002. A single domestication for maize shown by multilocus microsatellite genotyping. Proceedings of the National Academy of Sciences of the United States of America 99: 6080–6084.

54. McLaughlin CM, Li M, Perryman M, Heymans A, Schneider H, Lasky JR, Sawers RJH. 2024. Evidence that variation in root anatomy contributes to local adaptation in Mexican native maize. Evolutionary applications 17: e13673.

55. McSteen P. 2006. Branching out: the ramosa pathway and the evolution of grass inflorescence morphology. The plant cell 18: 518–522.

56. Mercer K, Martínez-Vásquez Á, Perales HR. 2008. Asymmetrical local adaptation of maize landraces along an altitudinal gradient. Evolutionary applications 1: 489–500.

57. Mercer KL, Perales H. 2019. Structure of local adaptation across the landscape: flowering time and fitness in Mexican maize (Zea mays L. subsp. mays) landraces. Genetic resources and crop evolution 66: 27–45.

58. Monier B, Bradbury P, Casstevens T, Jannink J-L, Buckler E. 2024. rPHG: R front-end for the practical haplotype graph.

59. Orr HA. 1998. Testing natural selection vs. genetic drift in phenotypic evolution using quantitative trait locus data. Genetics 149: 2099–2104.

60. Ortega-Paczka R. 2003. La diversidad del maíz en México. G. Esteva and C. Marielle, coordinators: 123–154.

61. Palaisa K, Morgante M, Tingey S, Rafalski A. 2004. Long-range patterns of diversity and linkage disequilibrium surrounding the maize Y1 gene are indicative of an asymmetric selective sweep. Proceedings of the National Academy of Sciences of the United States of America 101: 9885–9890.

62. Peiffer JA, Romay MC, Gore MA, Flint-Garcia SA, Zhang Z, Millard MJ, Gardner CAC, McMullen MD, Holland JB, Bradbury PJ, et al. 2014. The genetic architecture of maize height. Genetics 196: 1337–1356.

63. Perez-Limón S, Li M, Cintora-Martinez GC, Aguilar-Rangel MR, Salazar-Vidal MN, González-Segovia E, Blöcher-Juárez K, Guerrero-Zavala A, Barrales-Gamez B, Carcaño-Macias J, et al. 2022. A B73×Palomero Toluqueño mapping population reveals local adaptation in Mexican highland maize. G3 12.

64. Piperno DR, Moreno JE, Iriarte J, Holst I, Lachniet M, Jones JG, Ranere AJ, Castanzo R. 2007. Late Pleistocene and Holocene environmental history of the Iguala Valley, Central Balsas Watershed of Mexico. Proceedings of the National Academy of Sciences of the United States of America 104: 11874–11881.

65. Poplin R, Ruano-Rubio V, DePristo MA, Fennell TJ, Carneiro MO, Van der Auwera GA, Kling DE, Gauthier LD, Levy-Moonshine A, Roazen D, et al. 2018. Scaling accurate genetic variant discovery to tens of thousands of samples. bioRxiv: 201178.

66. Pya N, Wood SN. 2015. Shape constrained additive models. Statistics and computing 25: 543–559.

67. Pyhäjärvi T, Hufford MB, Mezmouk S, Ross-Ibarra J. 2013. Complex patterns of local adaptation in teosinte. Genome biology and evolution 5: 1594–1609.

68. R Core Team. 2022. R: A Language and Environment for Statistical Computing.

69. Reif JC, Warburton ML, Xia XC, Hoisington DA, Crossa J, Taba S, Muminović J, Bohn M, Frisch M, Melchinger AE. 2006. Grouping of accessions of Mexican races of maize revisited with SSR markers. TAG. Theoretical and applied genetics. Theoretische und angewandte Genetik 113: 177–185.

70. Romero Navarro JA, Willcox M, Burgueño J, Romay C, Swarts K, Trachsel S, Preciado E, Terron A, Delgado HV, Vidal V, et al. 2017. A study of allelic diversity underlying flowering-time adaptation in maize landraces. Nature genetics 49: 476–480.

71. Ruiz Corral JA, Durán Puga N, Sánchez González J de J, Ron Parra J, González Eguiarte DR, Holland JB, Medina García G. 2008. Climatic adaptation and ecological descriptors of 42 Mexican maize races. Crop science 48: 1502–1512.

72. Ruíz Corral JA, Sánchez González JJ. 1998. Base temperature and heat unit requirement of 49 Mexican maize races. Bergamo (Italy): Istituto Sperimentale per la Cerealicoltura,.

73. Runcie DE, Qu J, Cheng H, Crawford L. 2021. MegaLMM: Mega-scale linear mixed models for genomic predictions with thousands of traits. Genome biology 22: 213.

74. Salvi S, Sponza G, Morgante M, Tomes D, Niu X, Fengler KA, Meeley R, Ananiev EV, Svitashev S, Bruggemann E, et al. 2007. Conserved noncoding genomic sequences associated with a flowering-time quantitative trait locus in maize. Proceedings of the National Academy of Sciences of the United States of America 104: 11376–11381.

75. Sanchez G. JJ, Goodman MM, Stuber CW. 2000. Isozymatic and morphological diversity in the races of maize of Mexico. Economic botany 54: 43–59.

76. Savolainen O, Lascoux M, Merilä J. 2013. Ecological genomics of local adaptation. Nature reviews. Genetics 14: 807–820.

77. Scott MF, Ladejobi O, Amer S, Bentley AR, Biernaskie J, Boden SA, Clark M, Dell’Acqua M, Dixon LE, Filippi CV, et al. 2020. Multi-parent populations in crops: a toolbox integrating genomics and genetic mapping with breeding. Heredity 125: 396–416.

78. Shangguan W, Dai Y, Duan Q, Liu B, Yuan H. 2014. A global soil data set for earth system modeling. Journal of advances in modeling earth systems 6: 249–263.

79. Sigmon B, Vollbrecht E. 2010. Evidence of selection at the ramosa1 locus during maize domestication. Molecular ecology 19: 1296–1311.

80. Song B, Marco-Sola S, Moreto M, Johnson L, Buckler ES, Stitzer MC. 2022. AnchorWave: Sensitive alignment of genomes with high sequence diversity, extensive structural polymorphism, and whole-genome duplication. Proceedings of the National Academy of Sciences of the United States of America 119: e2113075119.

81. South A. 2011. rworldmap: A New R package for Mapping Global Data. The R Journal 3: 35–43.

82. Stinchcombe JR, Weinig C, Ungerer M, Olsen KM, Mays C, Halldorsdottir SS, Purugganan MD, Schmitt J. 2004. A latitudinal cline in flowering time in Arabidopsis thaliana modulated by the flowering time gene FRIGIDA. Proceedings of the National Academy of Sciences of the United States of America 101: 4712–4717.

83. Storey JD, Bass AJ, Dabney A, Robinson D. 2025. qvalue: Q-value estimation for false discovery rate control.

84. Strable J, Unger-Wallace E, Aragón Raygoza A, Briggs S, Vollbrecht E. 2023. Interspecies transfer of RAMOSA1 orthologs and promoter cis sequences impacts maize inflorescence architecture. Plant physiology 191: 1084–1101.

85. Twyford AD, Friedman J. 2015. Adaptive divergence in the monkey flower Mimulus guttatus is maintained by a chromosomal inversion. Evolution; international journal of organic evolution 69: 1476–1486.

86. Venkatesh TV, Harrigan GG, Perez T, Flint-Garcia S. 2015. Compositional assessments of key maize populations: B73 hybrids of the Nested Association Mapping founder lines and diverse landrace inbred lines. Journal of agricultural and food chemistry 63: 5282–5295.

87. Vielle-Calzada J-P, Martínez de la Vega O, Hernández-Guzmán G, Ibarra-Laclette E, Alvarez-Mejía C, Vega-Arreguín JC, Jiménez-Moraila B, Fernández-Cortés A, Corona-Armenta G, Herrera-Estrella L, et al. 2009. The Palomero genome suggests metal effects on domestication. Science 326: 1078.

88. Vollbrecht E, Springer PS, Goh L, Buckler ES 4th, Martienssen R. 2005. Architecture of floral branch systems in maize and related grasses. Nature 436: 1119–1126.

89. Wang L, Josephs EB, Lee KM, Roberts LM, Rellán-Álvarez R, Ross-Ibarra J, Hufford MB. 2021. Molecular Parallelism Underlies Convergent Highland Adaptation of Maize Landraces. Molecular biology and evolution.

90. Wellhausen EJ, Roberts LM, Hernandez-X. H. 1951. Razas de Maíz en México, su origen, características y distribución (PC Mangelsdorf, Ed.). Secretaría de Agricultura y Ganadería de México D. F.; Fundación Rockefeller.

91. Wellhausen EJ, Roberts LM, Hernandez-Xoconostle E, in collaboration with Mangelsdorf PC. 1952. Races of Maize in Mexico: their origin, characteristics and distribution. Jamaica Plain: Bussey Institution of Harvard University.

92. Wickham H. 2016. ggplot2: Elegant Graphics for Data Analysis.

93. Xu G, Wang X, Huang C, Xu D, Li D, Tian J, Chen Q, Wang C, Liang Y, Wu Y, et al. 2017. Complex genetic architecture underlies maize tassel domestication. The New phytologist 214: 852–864.

94. Yang Q, Li Z, Li W, Ku L, Wang C, Ye J, Li K, Yang N, Li Y, Zhong T, et al. 2013. CACTA-like transposable element in ZmCCT attenuated photoperiod sensitivity and accelerated the postdomestication spread of maize. Proceedings of the National Academy of Sciences of the United States of America 110: 16969–16974.

