## Supplemental Text and Figures for "The MexMAGIC population reveals the genetic architecture of clinal trait variation in Mexican native maize"

**Perez-Limón et al.**

### Supplemental Material

### Selection of MexMAGIC parents of analysis of source environments

*MexMAGIC parents*

| Founder | Short_Name | Accession | Type |
|---|---|---|---|
| Gordo | GOR | CHIH140 | OPV |
| Jala | JAL | NAYA6 | OPV |
| Mushito | MUS | MICH320 | OPV |
| Nal Tel | NAL | YUCA7 | OPV |
| Palomero Toluqueño | PAT | MEXI5 | OPV |
| Reventador | REV | Ames 30532 (MR18) | S6 |
| Tabloncillo | TAB | Ames 32914 (MR21) | S6 |
| Zapalote Chico | ZAP | Ames 30536 (MR23) | S6 |

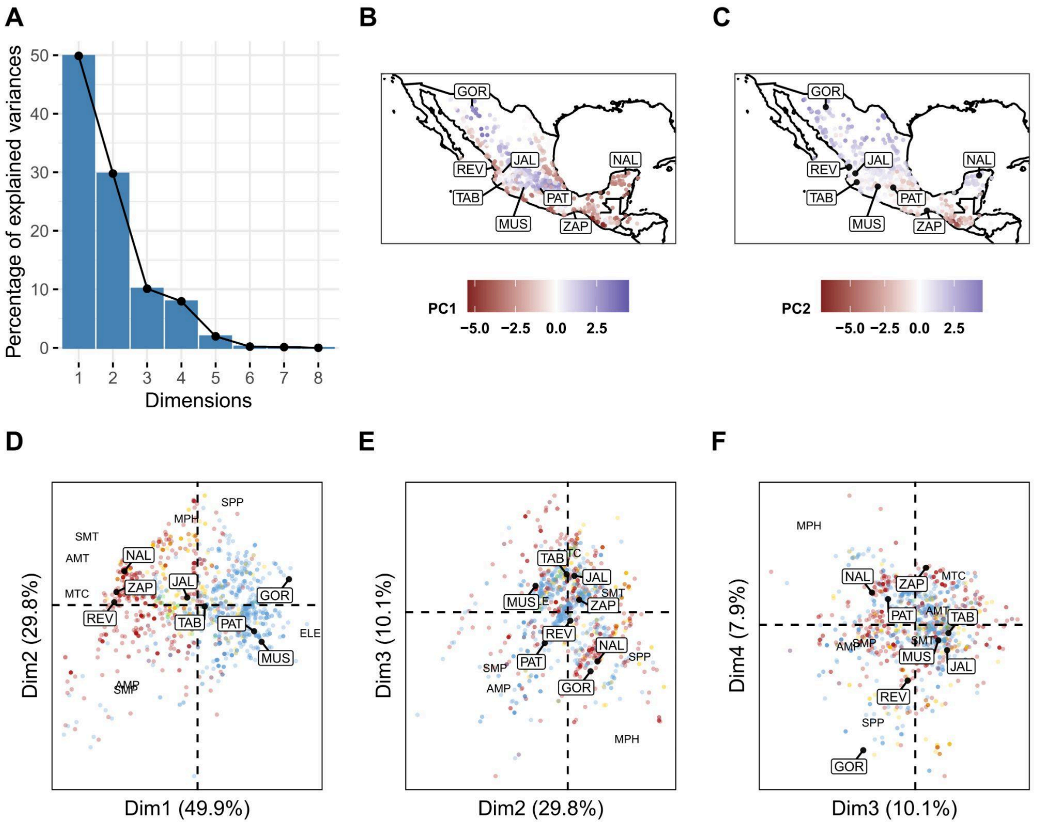

**Figures S1. Environmental Principal Component Analysis.**Principal Component (PC) analysis of the Mexican diversity panel and MAGIC founders using eight environmental descriptors and elevation as detailed in the main text. **A.** Scree plot showing the variance explained by the first eight PCs. **B., C.** Maps of source location for each individual colored by loading on PC1 and PC2, respectively. MAGIC founders labeled. **D., E., F.** PC biplots for the PCs (Dim) shown. Labels and coloring on individual points as main text Figure 1.

**Environmental clustering of Mexican diversity panel and MAGIC founders**

The MexMAGIC founders and Mexico diversity panel accessions were clustered by eight environmental descriptors (elevation, seasonal photoperiod, annual mean temperature, seasonal mean temperature, annual mean precipitation, seasonal mean precipitation, mean temperature of the coldest month and mean soil pH) using Partitioning Around Medoids (PAM) as previously described (Klein et al. 2020). Briefly, environmental descriptors were centered and scaled using R/caret (Kuhn 2021) and outlying values (> 3 standard deviations from the mean) removed. Within cluster sums of squares (WSS) were visualized as a function of cluster number using R/factoextra::fviz_nbclust (Kassambara and Mundt 2020). From inspection of the resulting curve, the accessions were grouped into four clusters using R/cluster::pam (Maechler et al. 2021) under default settings.

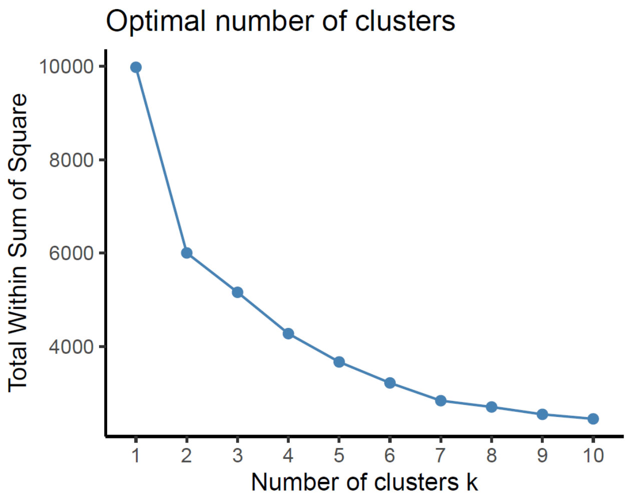


**Figures S2. Optimal number of environmental clusters.** Within cluster sum of squares as a function of cluster number for the MexMAGIC founders and Mexico diversity panel accessions clustered using eight environmental descriptors. Four was selected as the optimum number of clusters based on the “elbow” between 2 and 6 clusters.

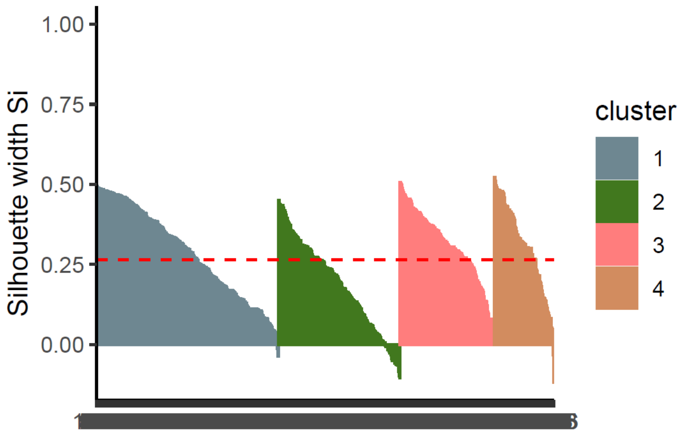


**Figures S3. Environmental cluster silhouette plot.** Within

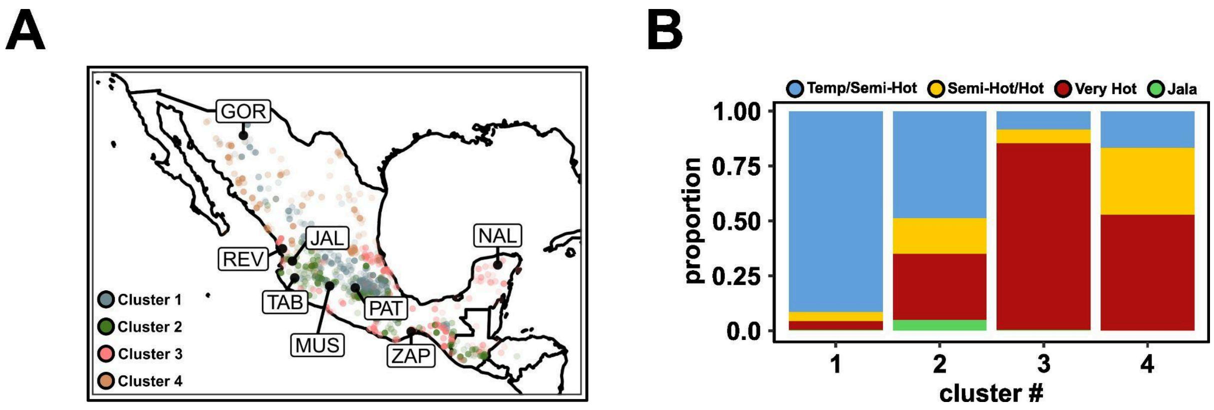


**Figure S4. Environmental clustering of Mexican diversity panel and MexMAGIC founders. A.** Source of the Mexican diversity panel labeled by environmental cluster assignment. The source of the eight MexMAGIC founder accessions is shown by boxed three-letter identifiers. **B.** Composition of environmental clusters with respect to previously assigned climate groups (Ruiz Corral et al. 2008).

**Crossing scheme for MAGIC families**

*F1 Generation*

Valle de Banderas, Nayarit, Mexico, Winter 2015/2016

A single ear was advanced for each of four F1 combinations, capturing the eight target parents.

| F1 ear | Mum_stock | Dad_stock | Mum_geno | Dad_geno |
|---|---|---|---|---|
| A | RS16-701.1 | RS16-344.4 | TAB | MUS |
| B | RS16-141.10 | RS16-1032.6 | JAL | PAT |
| C | RS16-1033.8 | RS16-144.9 | GOR | REV |
| D | RS16-341.2 | RS16-350.2 | NAL | ZAP |

*Four-way Cross*

Irapuato, Guanajuato, Mexico, Summer 2016.

A single $F_1$ plant was selected for each $F_1$ combination and used as both male and female in distinct four-way crosses, ensuring that only a single haplotype was captured per founder parent. The four plants representing the $F_1$ crosses A, B , C and D, were crossed in a chain as AxB, BxD, DxC and CxA. 120 seeds were advanced from each four-way ear, reflecting the lowest yielding cross. The crosses BxD and CxA were designated A1 and A2, respectively, capturing a complete set of the eight founder parents. Similarly, the crosses DxC and AxB, were designated B1 and B2.

| 4-way ear | Mum_stock | Dad_stock | Mum_geno | Dad_geno |
|---|---|---|---|---|
| A1 | RS16-9538.1 | RS16-9548.4 | B: JALxPAT | D: NALxZAP |
| A2 | RS16-9547.2 | RS16-9545.4 | C: GORxREV | A: TABxMUS |
| B1 | RS16-9548.4 | RS16-9547.2 | D: NALxZAP | C: GORxREV |
| B2 | RS16-9545.4 | RS16-9538.1 | A: TABxMUS | B: JALxPAT |

*Eight-way Cross*

Valle de Banderas, Nayarit, Mexico, Winter 2016/2017.

All 120 seeds were planted from each of the four four-way ears A1, A2, B1 and B2. A1 individuals were used to pollinate A2 mothers and vice versa, although an effort was made to avoid reciprocal crosses between any two individuals. Similarly, crosses were made between B1 and B2 individuals. Eighty ears were advanced from each of the A1/A2 and B1/B2 intercrosses, for a total of 160 families.

*Intermating 1*

Valle de Banderas, Nayarit, Mexico, Winter 2017/2018.

Each eight-way ear was planted as a row for 80 A rows (progeny of eight-way A1xA2) and 80 B rows (progeny of eight-way B1xB2). As many pollinations as possible were made between A and B rows. A and B rows were used as both male and female, but no single AxB combination was made more than once. 500 ears were advanced.

*Intermating 2*

Irapuato, Guanajuato, Mexico, Summer 2018.

Five replicate single-seed bulks were made from the 500 ears recovered from Intermating 1. Individuals were intermated within bulks. A total of 474 ears were recovered across all bulks.

*Intermating 3*

Valle de Banderas, Nayarit, Mexico, Winter 2018/2019.

Three replicate single-seed bulks were made from the 474 ears recovered from Intermating 2. Individuals were intermated within bulks. A total of 390 ears were recovered across all bulks.

*Selfing 1 to 3*

Ameca, Jalisco, Mexico Summer 2019; San Juan de Abajo, Nayarit, Mexico Winter 2019/2020; Ameca, Jalisco, Mexico Summer 2020.

The replicate single-seed bulks were made from the 390 ears recovered from Intermating 3. Individuals were self-pollinated in Summer 2019 to recover 328 S1 families. These were advanced as ear-to-row through two more generations to generate S3 families MEMA001 to MEMA328. Not all families were recovered in any given planting, and additional nurseries were used in Valle de Banderas, Nayarit, Mexico, Winters 2021/22 and 2022/2023, and Guadalajara. Jalisco, Mexico, Summer 2023 to complete the S generations. Additional families were derived from S1 ears produced 1) in Irapuato, Guanajuato, Mexico, Summer 2019 from an additional 390 seed I3 bulk (MEMA329 to MEMA376), 2) from a fourth intermating of I3 bulks in Valle de Banderas, Nayarit, Mexico, Winter 2019/2020 with subsequent self-pollination in the same location Winter 2021/22 (MEMA377 to MEMA474), and 3) from I3 single seed bulk seed originally selected for small (MEMA475 to MEMA514; MEMA555 to MEMA594) or large (MEMA515 to MEMA554; MEMA595 to MEMA633) seed size. A total of 609 families were advanced to the S3 generation.

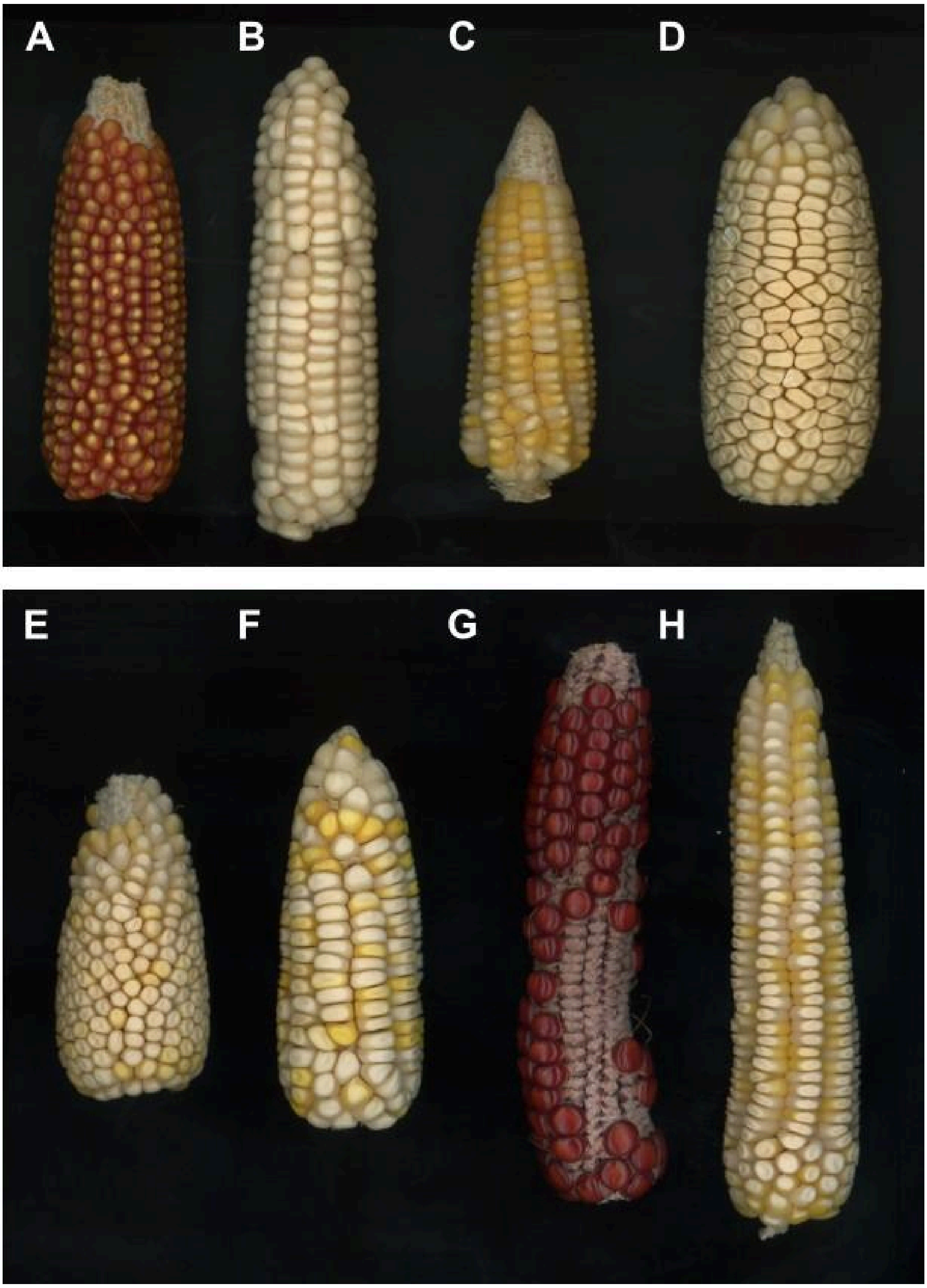


**Figures S5. Early generation ears. A - D. Example $F_1$ ears**. **A.** Oloton x Mushito. **B.** Gordo x Reventador. **C.** Nal Tel x Tabloncillo. **D.** Jala x Palomero Toluqueño. Ear D was used in the final population. Ears B and C are representative of Gordo and Nal Tel varieties, respectively, although these were incorporated into the final population as the crosses Gordo x Reventador and Nal Tel x Zapalote Chico. **E-G. Four-way ears. E.** 'A1' (Jala x Palomero Toluqueño) x (Nal Tel x Zapalote Chico). **F.** 'B1' (Nal Tel x Zapalote Chico) x (Gordo x Reventador). **G.** 'A2' (Gordo x Reventador) x (Tabloncillo x Mushito). **H.** 'B2' (Tabloncillo x Mushito) x (Jala x Palomero Toluqueño).

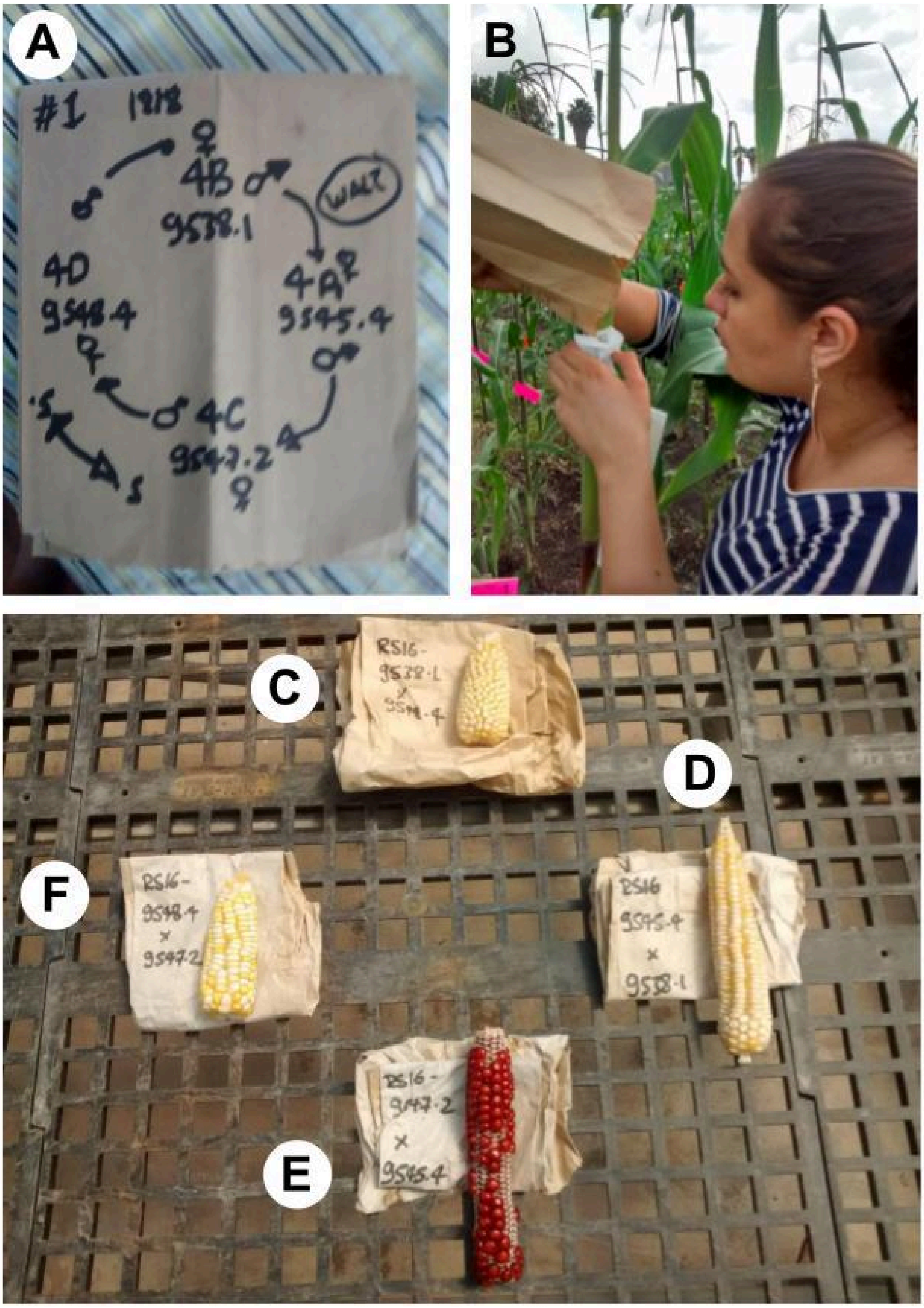


**Figures S6. Four-way crosses. A.** Crossing scheme using four F1 individuals in a closed chain as both male and female. **B.** Dr. G. Carolina Cintora Martinez performs the four-way crosses, Irapuato, Mexico, Summer 2016. **C.** 'A1' (Jala x Palomero Toluqueño) x (Nal Tel x Zapalote Chico). **D.** 'B2' (Tabloncillo x Mushito) x (Jala x Palomero Toluqueño). **E.** 'A2' (Gordo x Reventador) x (Tabloncillo x Mushito). **F.** 'B1' (Nal Tel x Zapalote Chico) x (Gordo x Reventador).

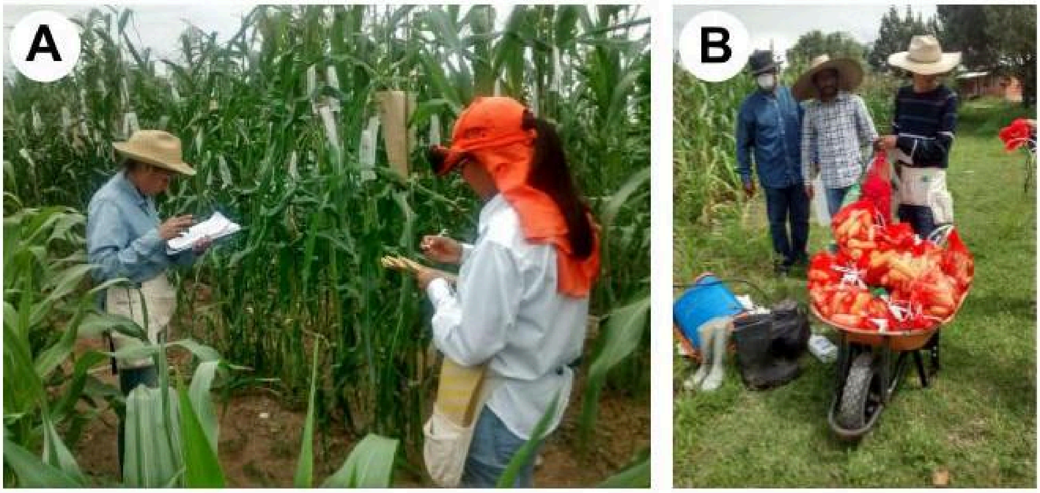


**Figures S7. Intermating Generations. A.** Intermating 1, Valle de Banderas, Nayarit, Mexico, Winter 2017/2018. Dr. M. Rosario Ramírez-Flores (left) and Dr. Ana Laura Alonso-Nieves (right). **B.** Harvesting Intermating 2, Irapuato, Mexico, Fall 2018.

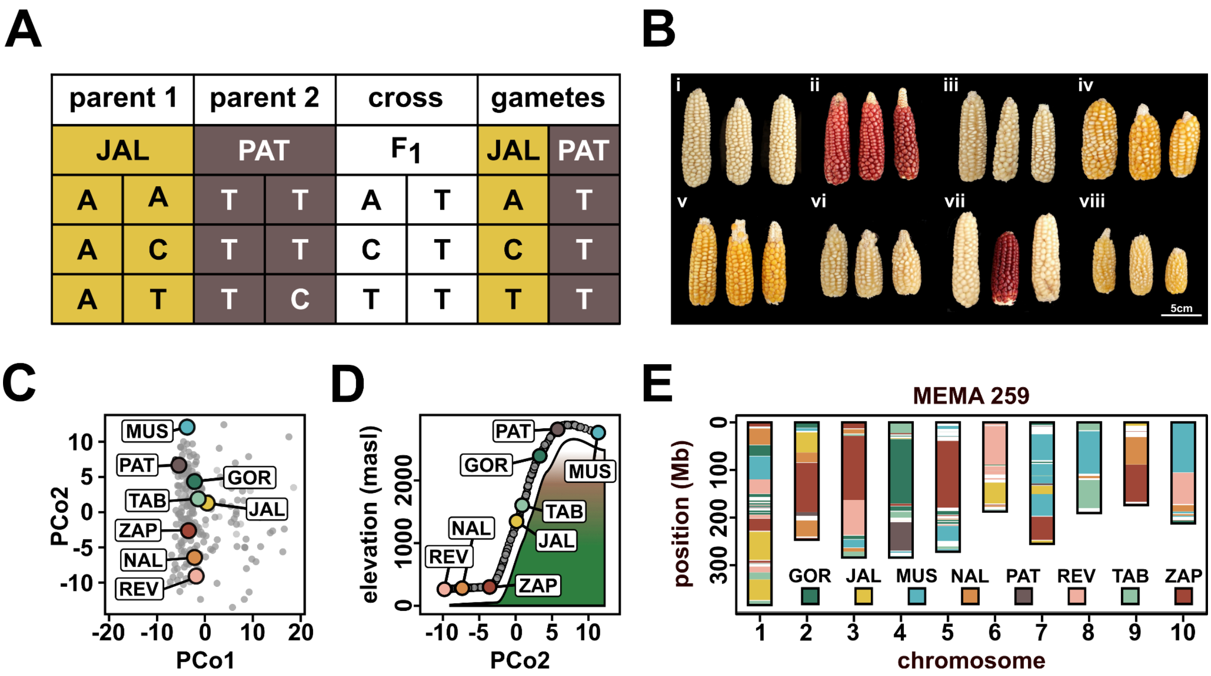


**Figure S8. Genetic characterization of the MexMAGIC families. A.** Triplet approach to inference of parental gametes in the $F_1$ cross. **B.** Bulked S3 ears of example MexMAGIC families. i) MEMA041, ii) MEMA314, iii) MEMA297, iv) MEMA045, v) MEMA315, vi) MEMA040, vii) MEMA042, viii) MEMA057. **C.** Genotypic Principal Coordinate (PCo) analysis of MEMA families and founder parents using a distance matrix based on 10,191 chip-called SNPs. **D.** LOESS fit (filled shape) of PCo2 against elevation for the founder parents (colored, labeled points). Grey points show a predicted elevation for the MEMA families based on the LOESS fit. Points are vertically displaced from the fitted line for visualization. **E.** Founder ancestry across the ten chromosomes of an example MEMA family.

Regions were assigned using an ancestry probability threshold of 0.9. Regions for which no individual parent passed the threshold were set as non-assigned and not colored here.

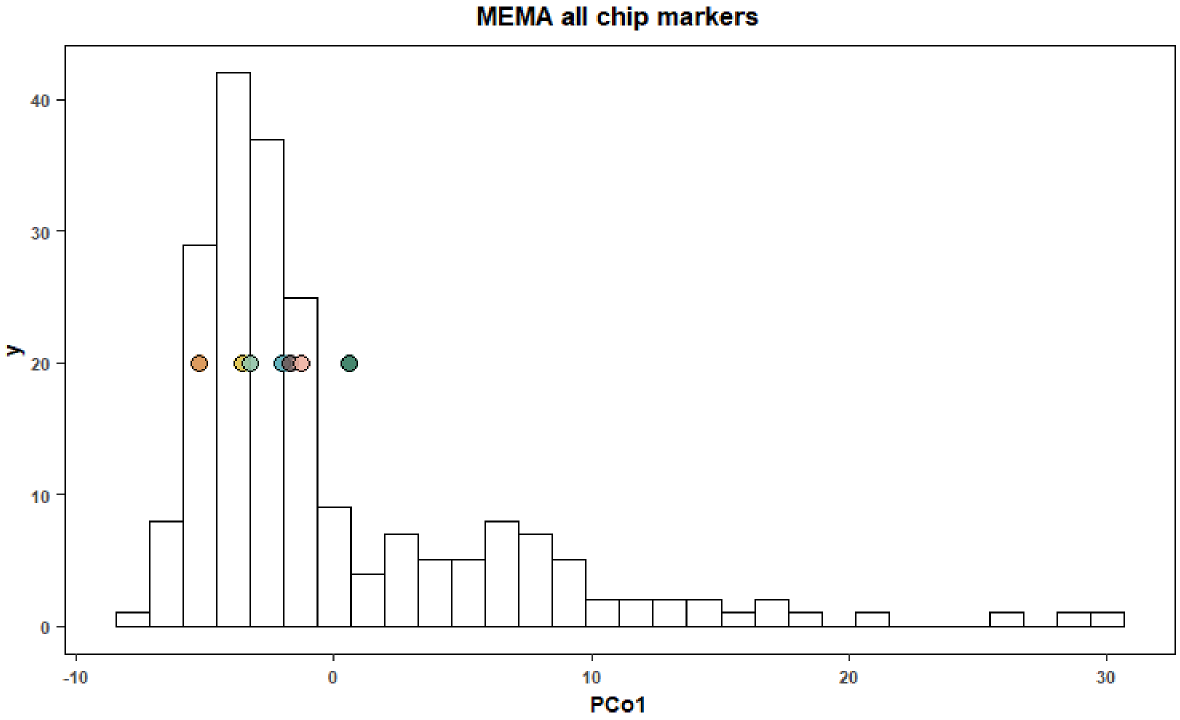


**Figure S9. Outlying families in genetic PCo1.** Genotypic Principal Coordinate (PCo) analysis of MEMA families and founder parents using a distance matrix based on 10,191 chip-called SNPs. Frequency of PCo1 loading across 200 families and 8 founders. Colored points show PCo1 loading of the eight founders, colors as main text figures.

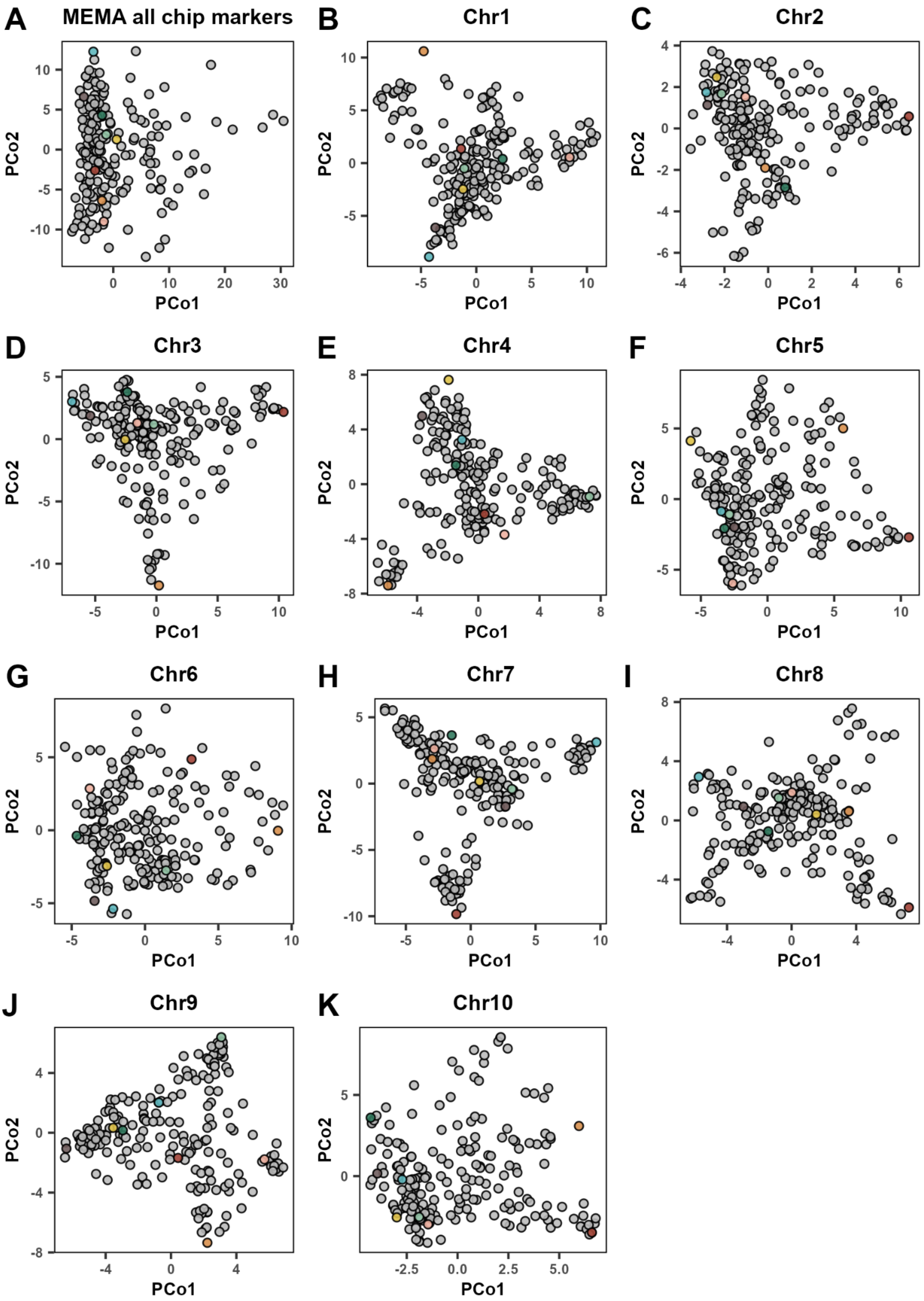


**Figure S10. Genetic characterization of the MexMAGIC families.** Genotypic Principal Coordinate (PCo) analysis of MEMA families and founder parents, run separately for each chromosome.

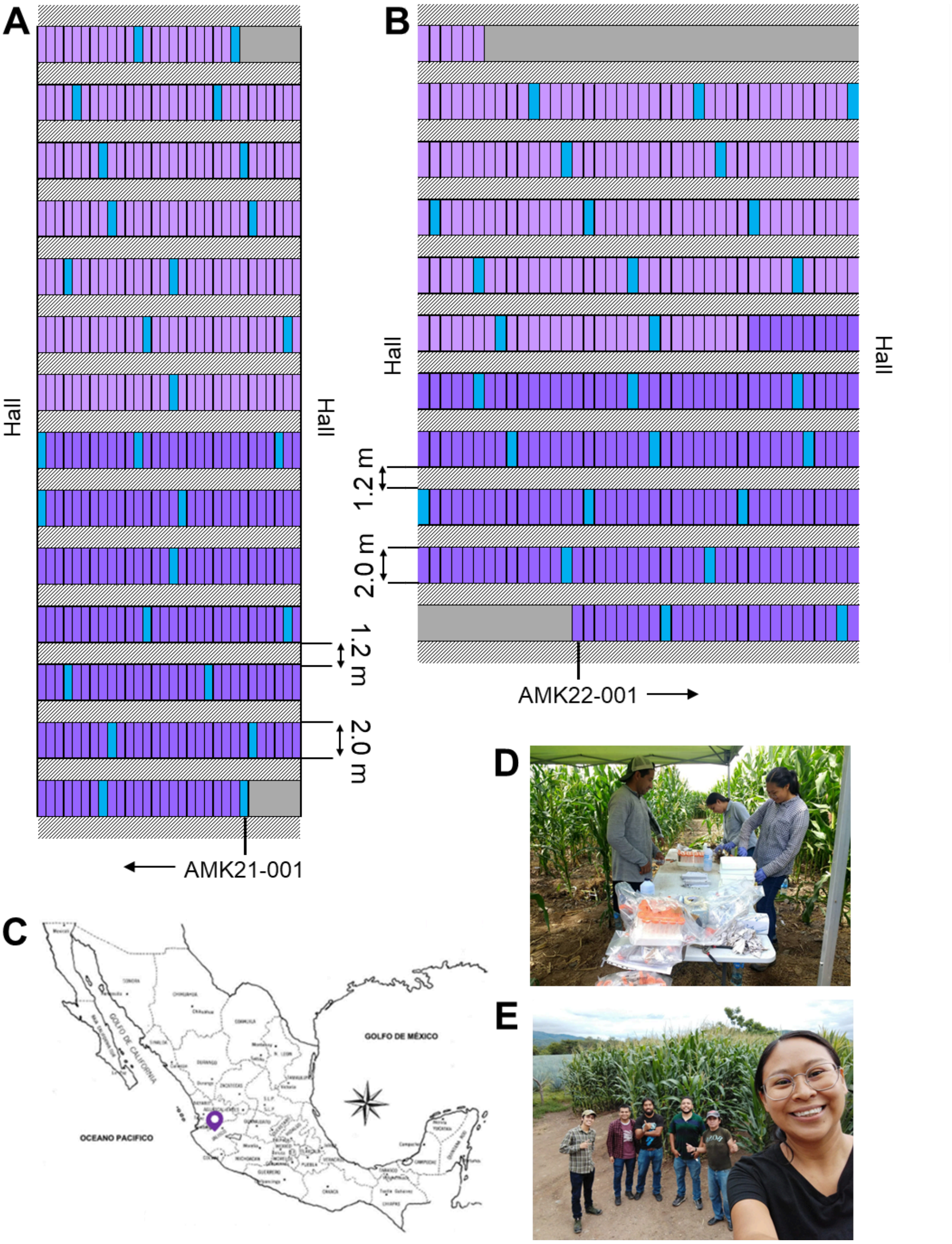


**Figure S11.** Field evaluation of the MexMAGIC testcross population (MexMAGIC_TC). Evaluations were performed in 2021 and 2021 A-B. Field design maps of the 2021 and 2022 evaluations, respectively. Families were planted randomly in two independent blocks. Purple color indicates block 1, and light purple indicates block 2. Blue color shows the distribution of the commercial hybrid used as check C. D-E. Teamwork during of 2021 and 2022.

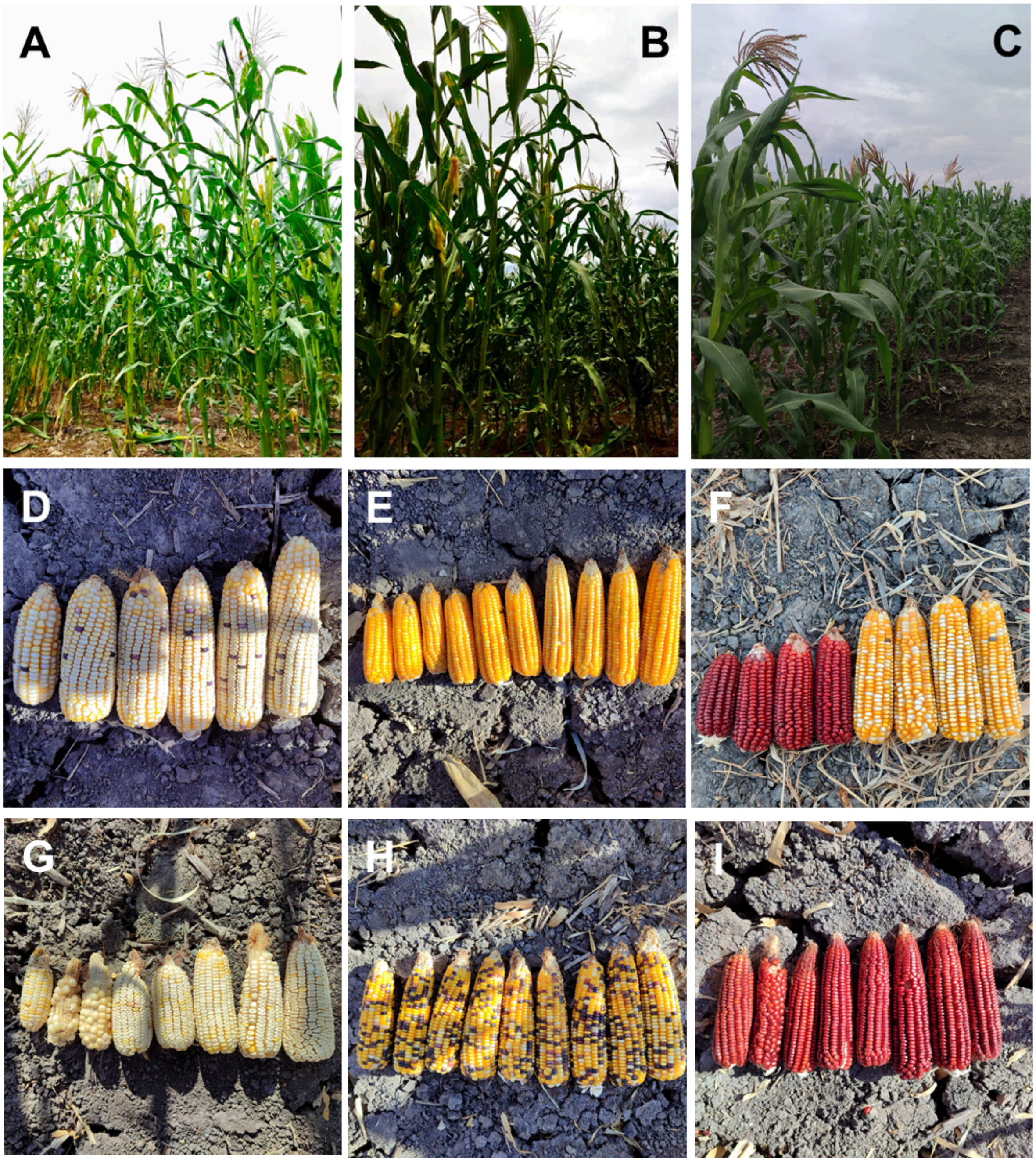


**Figure S12. Field evaluation Ameca 2022**

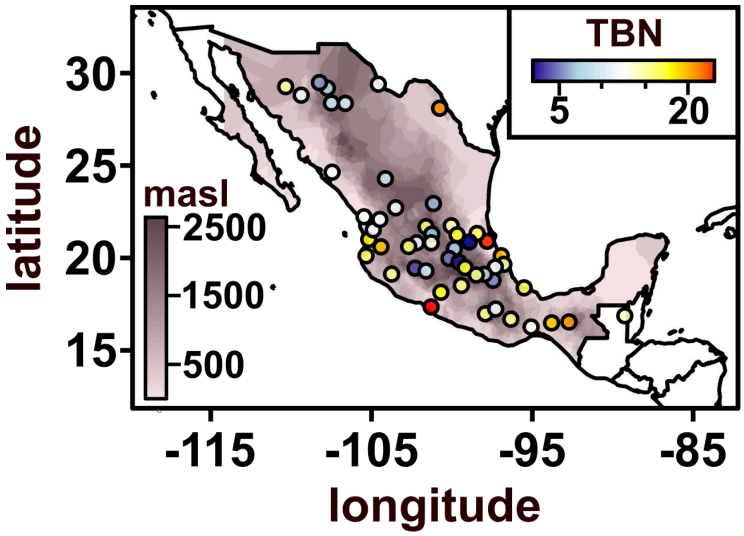


**Figure S13.** Sampling of Mexican native varieties reported in Janzen et al., 2022. Points shaded by tassel branch number (TBN) in common garden evaluation.

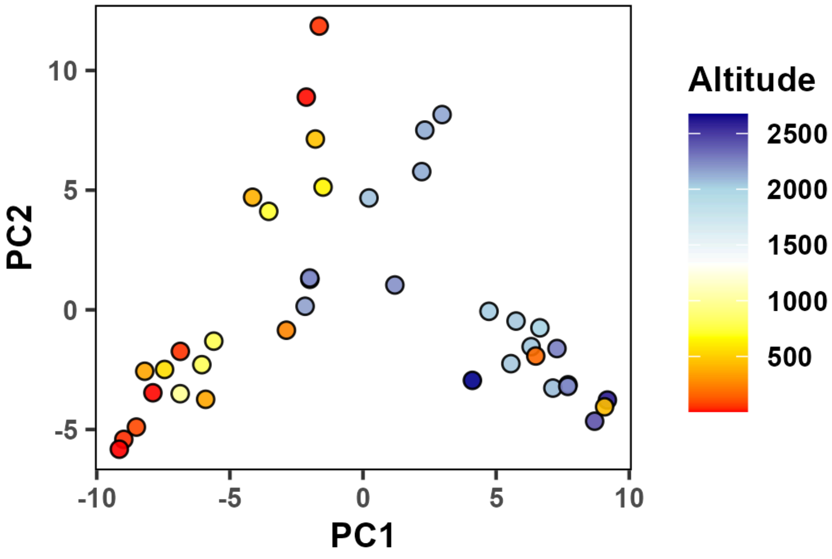


**Figure S14. Population structure in the Mexican HiLo panel.** Multidimensional scaling (principal coordinates) for 40 accessions (Janzen et al. 2022) using 4060 SNPs. The lowland accession (CIMMYTMA-020752) with high loading on PCo1 (PC1 = 9.07, elevation = 436 masl) is classified as Conico, a highland variety, and showed low tassel branch number (TBN = 1.75). CIMMYTMA-020752 was collected near La Reforma, Hidalgo (21.06234 N, -98.87007 W) in a broadly highland region. A second anomalous accession (CIMMYTMA-007076; PC1 = 6.49, elevation = 177 m) is classified as Bofo and was collected north of Tepic, Nayarit (21.71667 N, -104.88333 W).

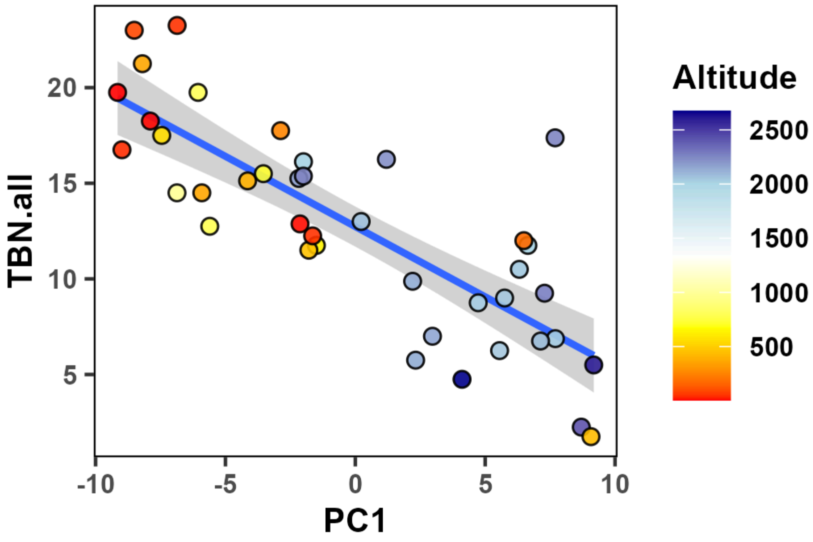


**Figure S15. The Mexican HiLo panel is strongly structured by elevation.** Population PCo1 (PC1) is strongly correlated ($p < 0.001$) with tassel branch number (TBN.all) in the HiLo panel. Line shows liner fit +/- SE.

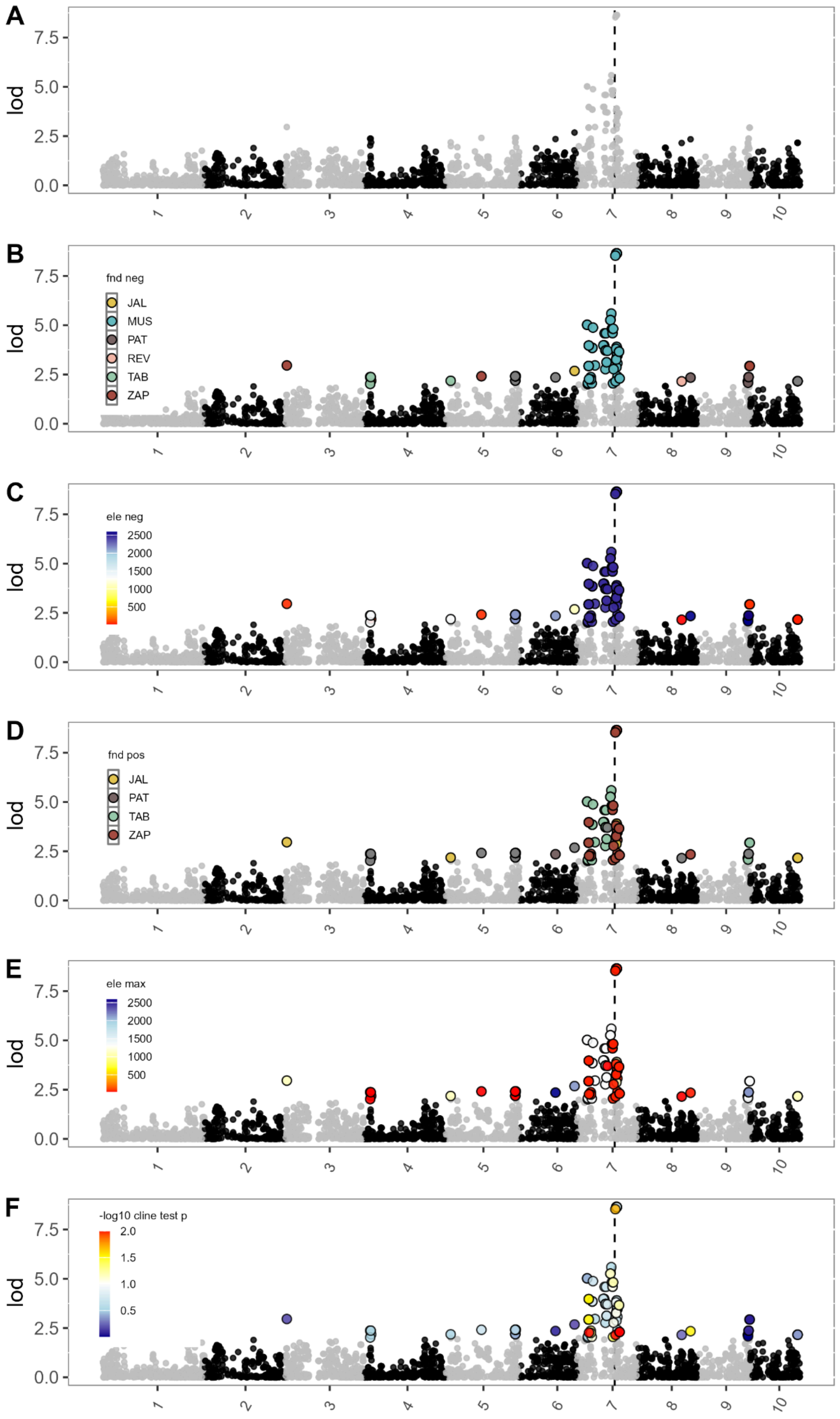
A
B
C
D
E
F
lod
7.5
5.0
2.5
0.0
1
2
3
4
5
6
7
8
9
10
fnd neg
JAL
MUS
PAT
REV
TAB
ZAP
ele neg
2500
2000
1500
1000
500
fnd pos
ele max
-log10 cline test p
2.0
1.5
1.0
0.5

**Figure S16.** GWAS support (LOD) in the MexMAGIC for association with TBN across the ten maize chromosomes. All SNPs with LOD > 2 are shown. A reduced random sampling of lower-supported SNPs is shown. The physical position of the candidate gene *Ra1* is indicated by a dashed vertical line. A) All SNPs shaded to show chromosome division. B-F) SNPs with LOD > 2 are colored by B) the founder showing the most negative effect on TBN at that SNP, C) the source elevation of the founder showing the most negative effect on TBN at that SNP, D) the founder showing the most positive effect on TBN at that SNP, E) the source elevation of the founder showing the most positive effect on TBN at that SNP, F) support for a second model fitting allele source elevation to TBN.

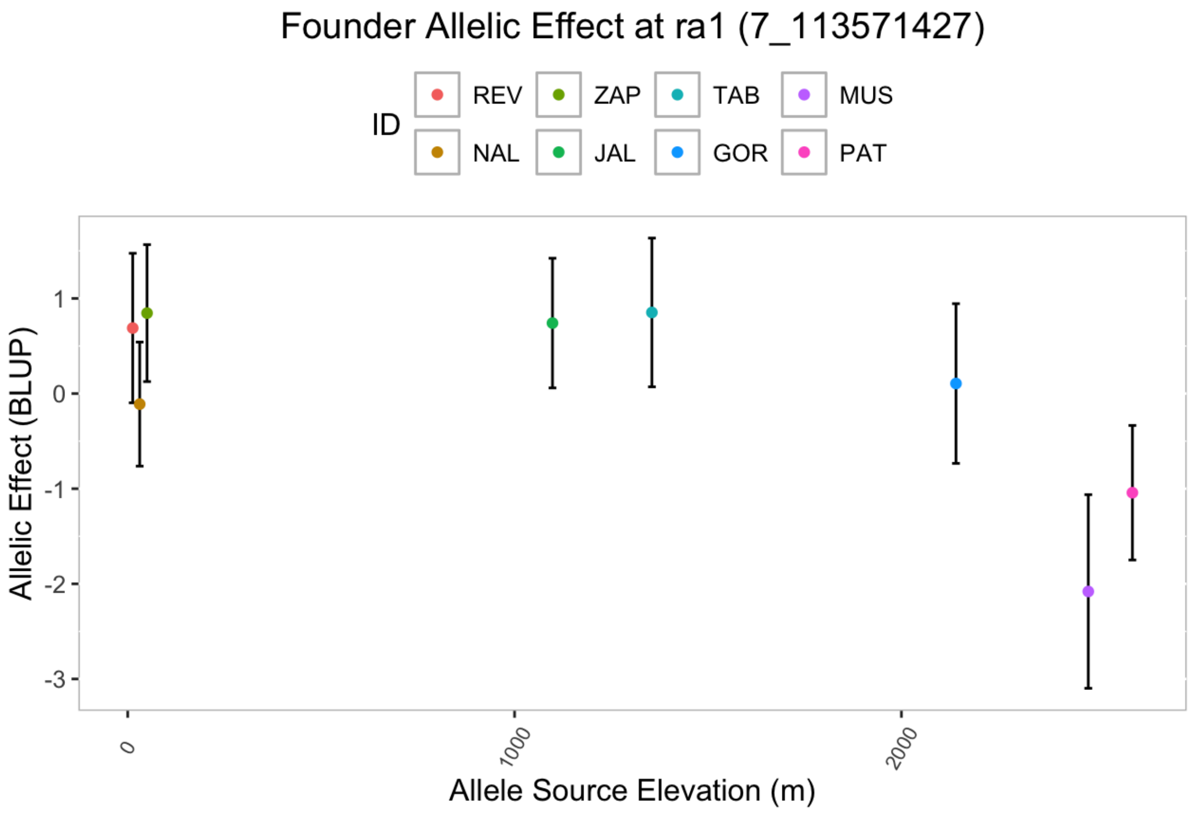


**Figure S17. Founder allelic effect *Ra1* (S7_113571427).** The x axis reflects the Allele source elevation in meters. The point represents the BLUP allelic effect and the error bars are +/- one standard error, estimated with the {R/qtl2::fit1} function. The color of the dots represents the founders of the MexMAGIC population.

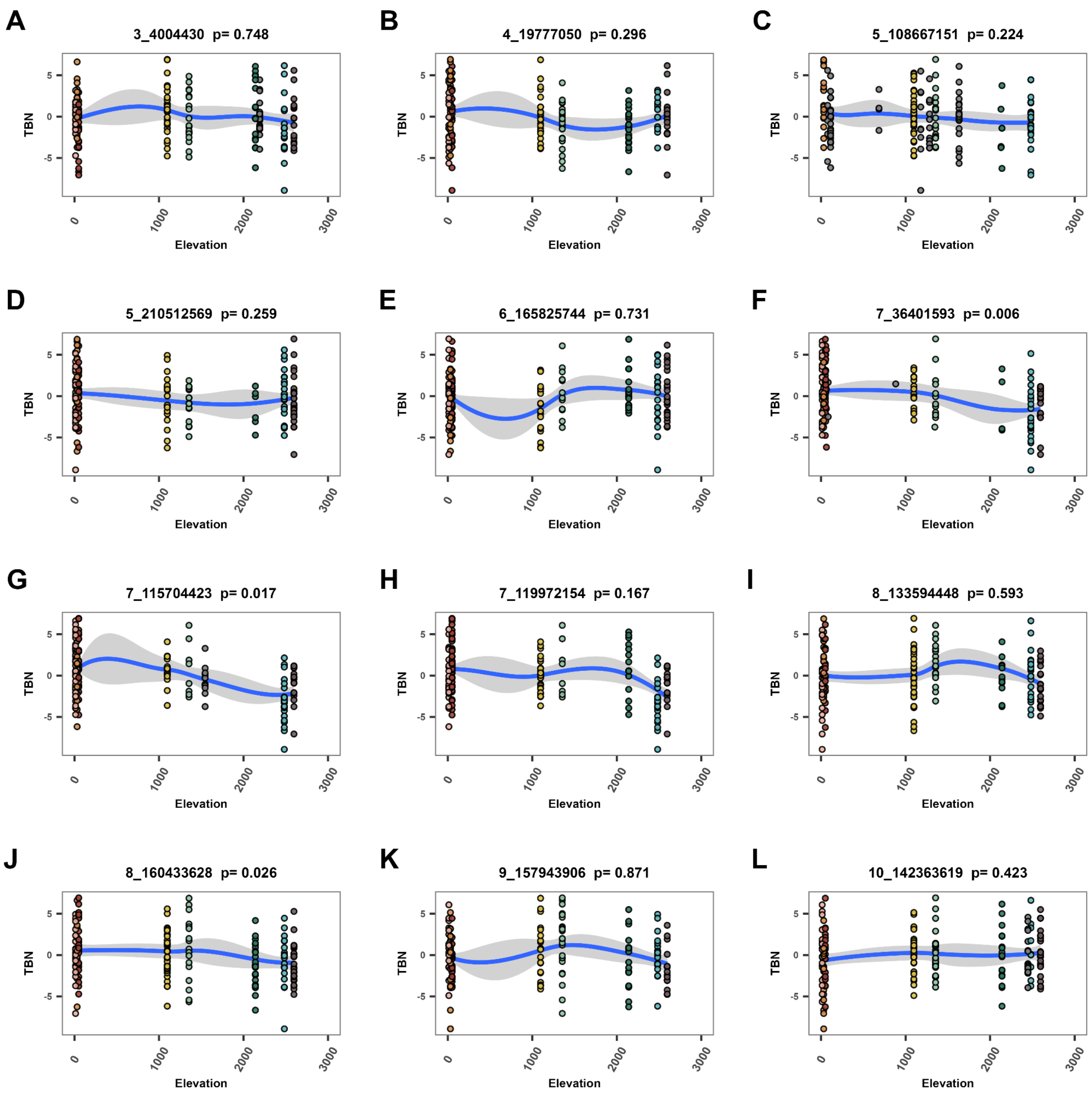


**Figure S18. Tassel branch number allele effects with respect to source elevation. A-L** Relationship between allele source elevation at the named marker and TBN for the MexMAGIC families. Source elevation was calculated as the product of genotype probabilities at each marker and the founder elevations. Points are colored by the founder hard-call at main text Fig.3. Uncolored points represent families for which the founder call was ambiguous. Blue line shows a LOESS fit, with the area extending +/- 1 standard error. Model statistics refer to a linear fit following the model described in the main text.

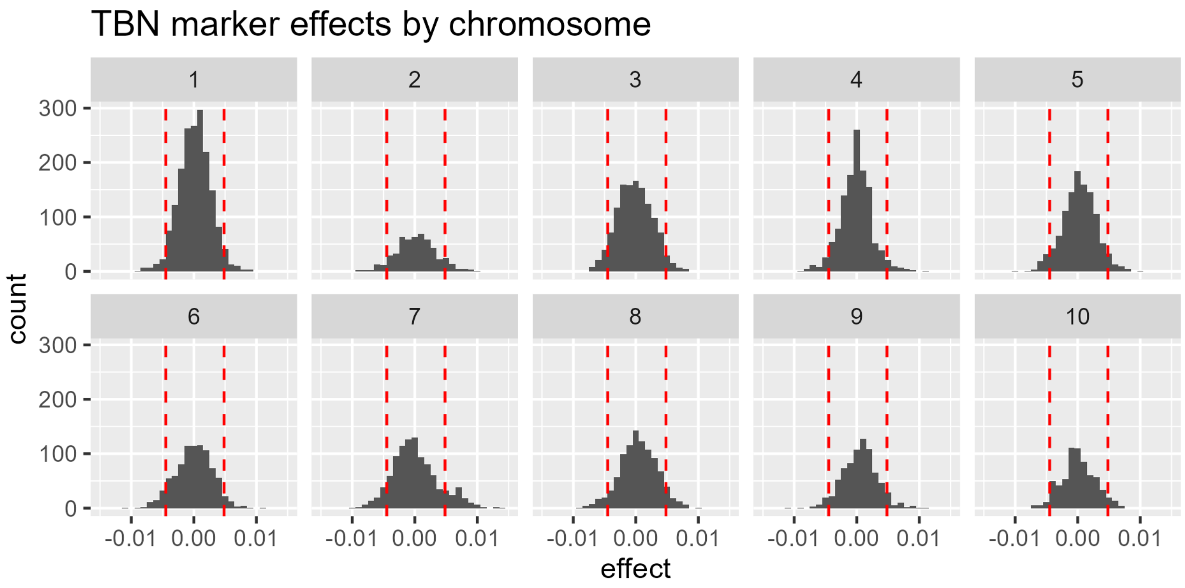


**Figure S19. Genomewide TBN marker effects.** Distribution of genomewide marker effects on Tassel Branch Number from the MexMAGIC. Red dotted lines represent +/- 2 SDs.

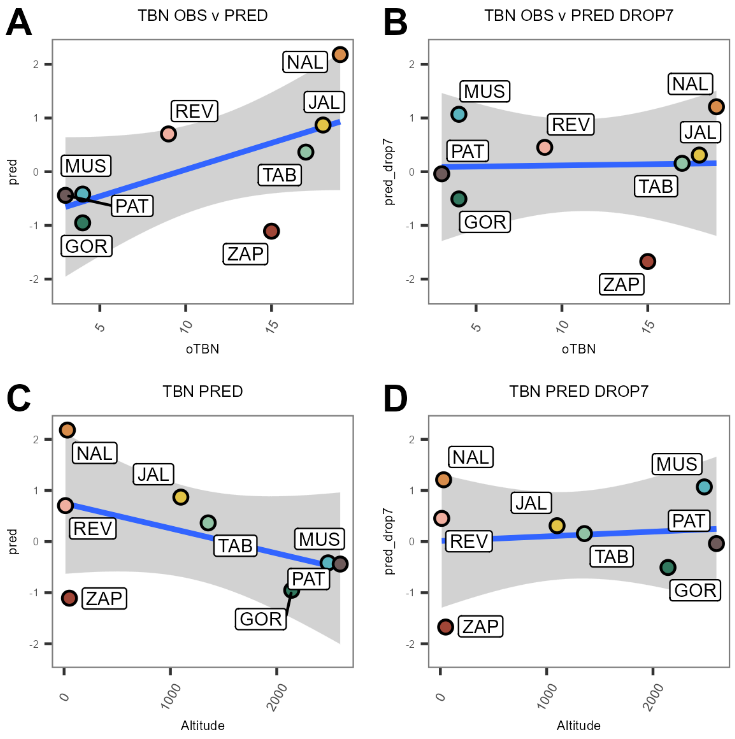


**Figure S20. Genomewide prediction of tassel branch number. A.** Founder Tassel branch number (TBN) predicted from genomewide marker effects in the MexMAGIC family against observed founder TBN. **B**. as A, dropping markers on chromosome 7, including the major QTL at *Ra1*. **C., D.** as A.,B., substituting founder elevation for observed founder TBN.

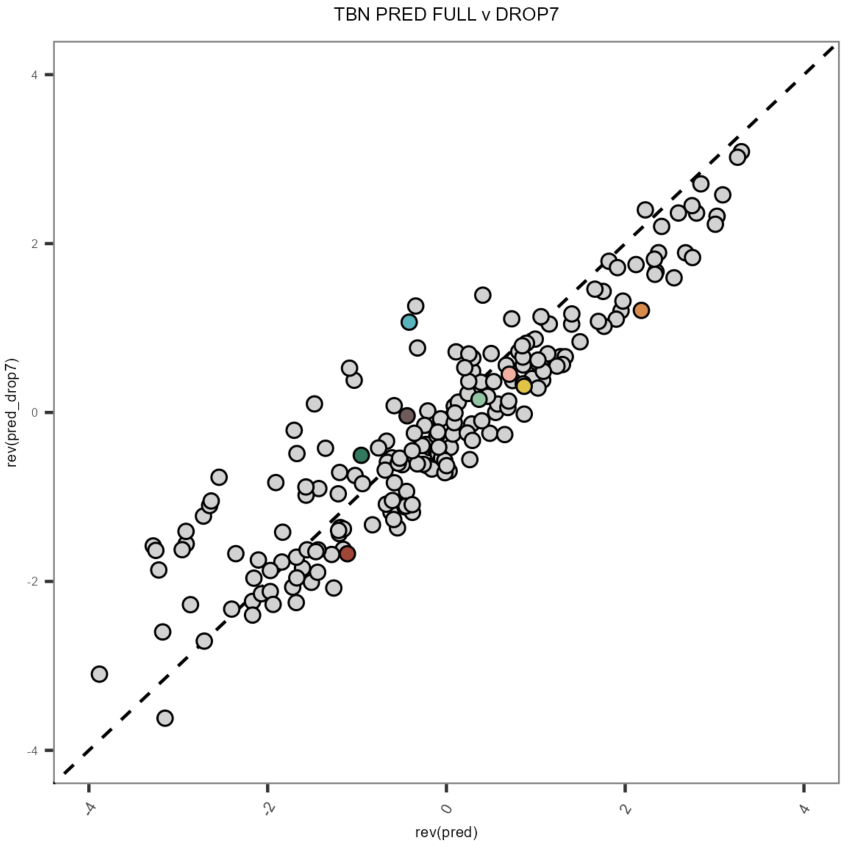


**Figure S21. Impact of dropping chromosome 7 markers on predicted TBN.** Predicted TBN (BLUP) for founders (colored as main text figures) and MexMAGIC families using a full genomewide markers (x-axis) or dropping markers on chromosome 7, including the major QTL at *Ra1* (y-axis).

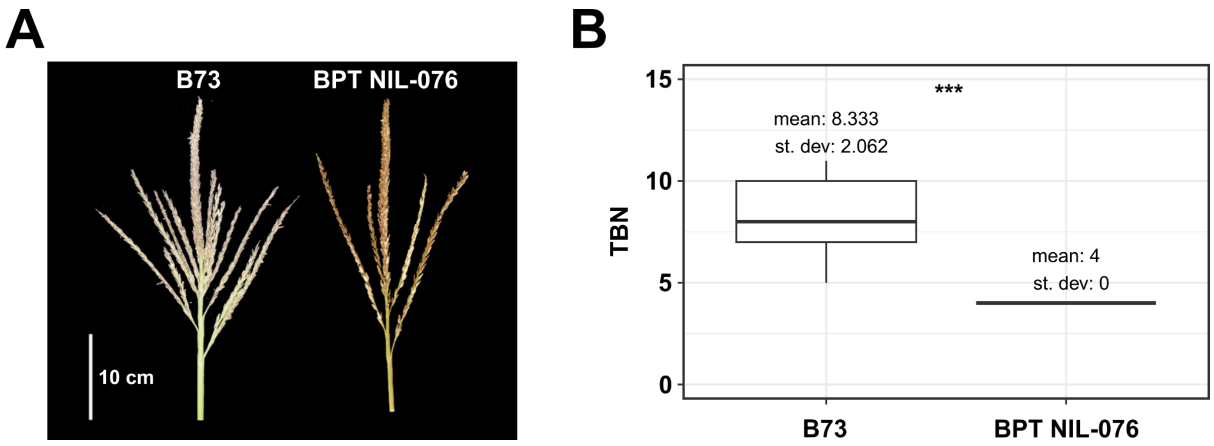


**Figure S22. Tassel branching in B73-*Ra1-PT* BC3DH NIL**. A) B73 tassel (left) and tassel of a BC3 NIL (BPT NIL-076) carrying the *Ra1* locus from Palomero Toluqueño in the B73 background. B) Tassel Branch number (TBN). n = 9 and 10 for B73 and the NIL, respectively.

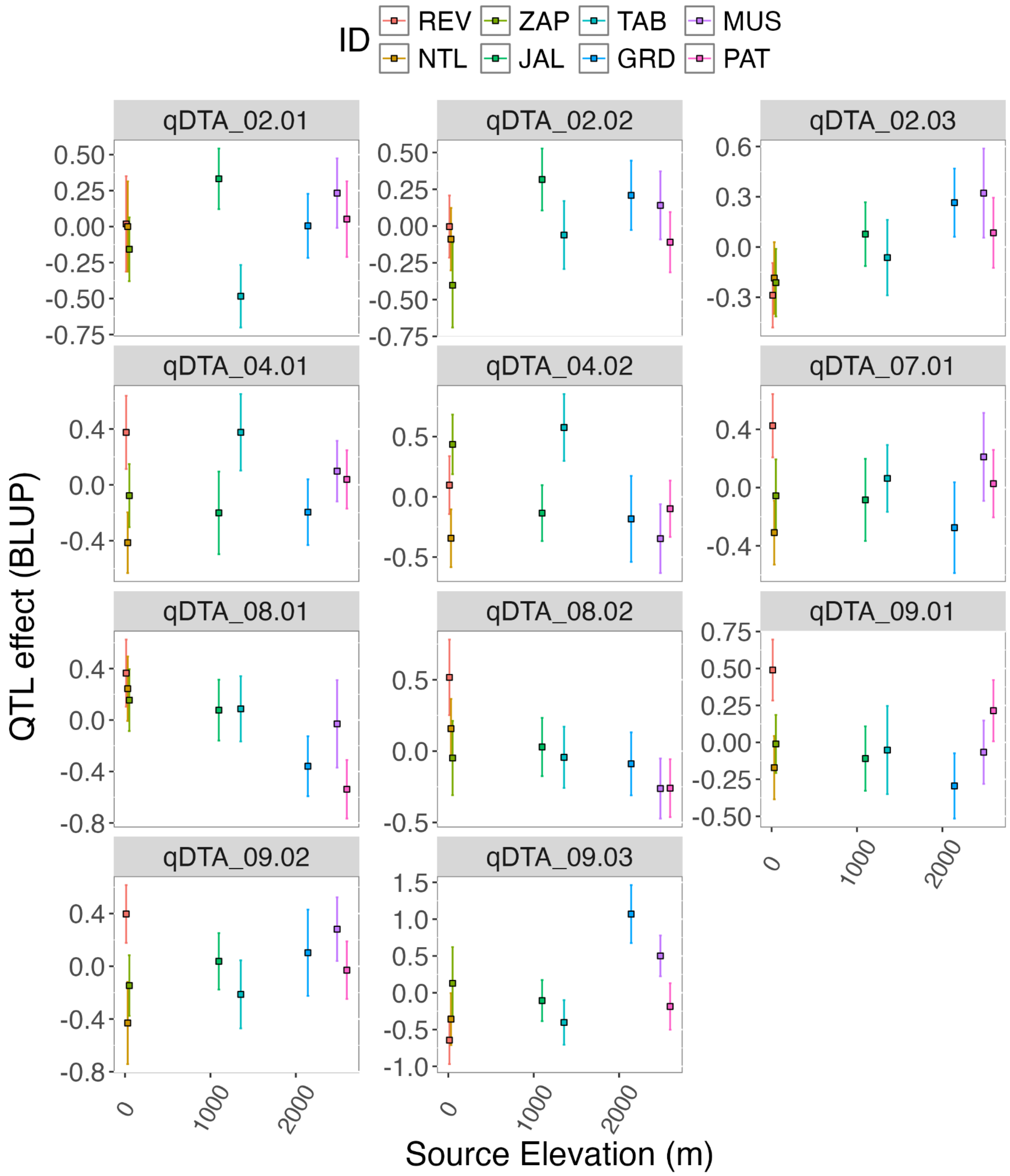


**Figure S23. Effect plots at the lead SNP for the 11 QTLs identified for DTA in the MexMAGIC population along the source altitude of the founder.** The x axis shows founder source elevation and the y axis the estimated allelic effect of the founder at the QTL (BLUP).

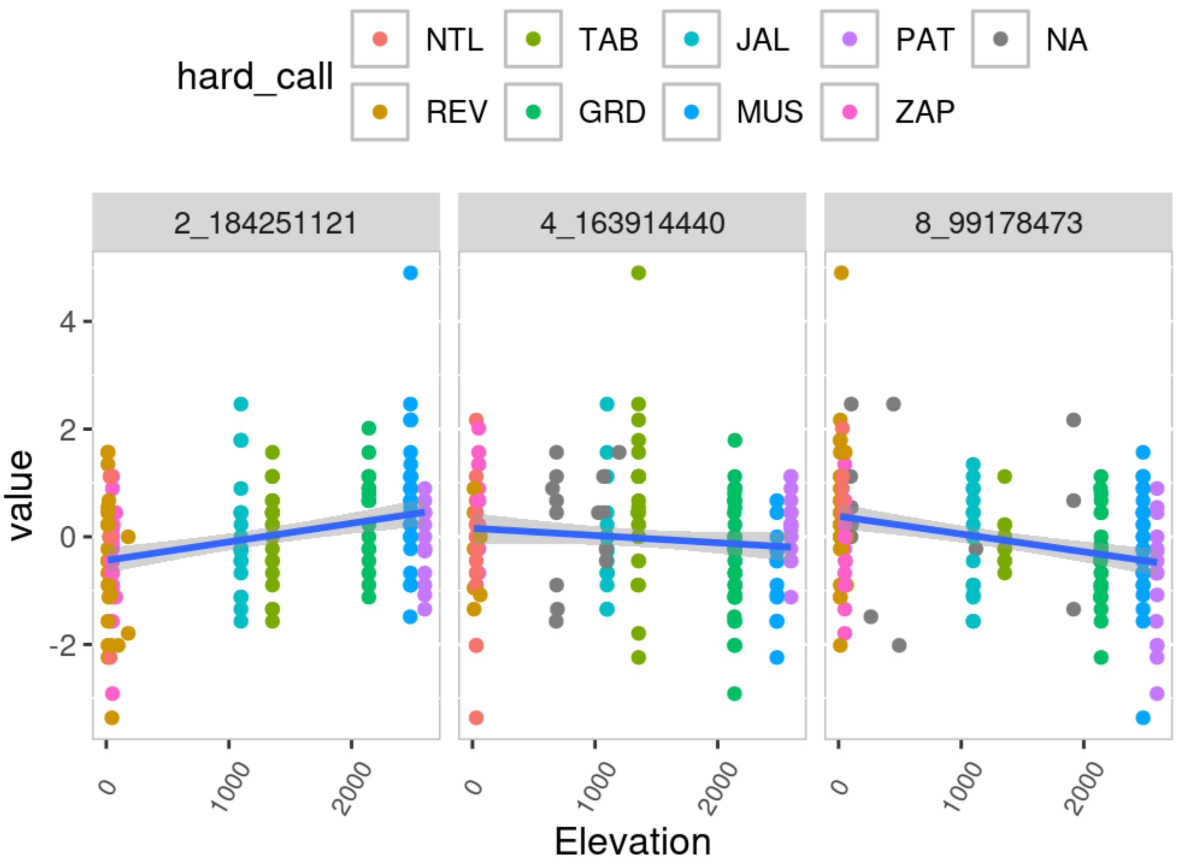


**Figure 24. Flowering time founder effect along an elevation cline.** X axis represents the source elevation, and Y axis represents the phenotypic value (BLUP). Each dot represents a family of the MexMAGIC population and the color of the founder genotype at each marker (genotype probability >= 0.9); families whose greatest founder probability was < 0.9, were considered as NA. The line represents a linear fit, and the shading the confidence interval around the linear fit.

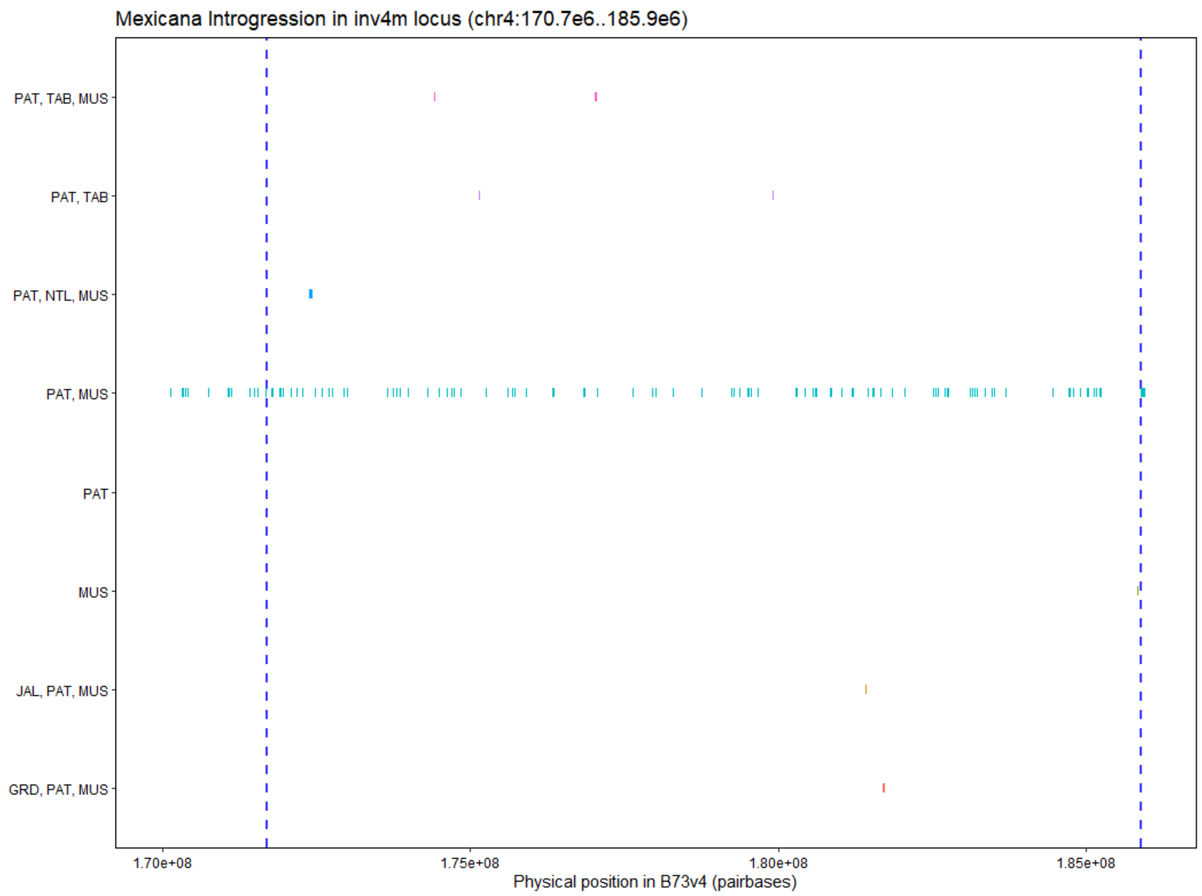


**Figure S25. *Zea mays* sp. *mexicana* introgression at *Inv4m*.** *Zea mays* sp. *mexicana* introgression tracks detected in the MexMAGIC founder genomes around the *inv4m* region (chromosome 4, ~ 170 - 186 MC) using ELAI (Guan 2014) according to methods in (Yang et al. 2023). Adjacent SNPs with an ELAI score >= 1.8 were collapsed into tracks using the {R/GenomicRanges} package (Lawrence et al. 2013). The vertical colored lines represent the tracks detected for a single or any combination of founders. Consistent evidence for introgression along *inv4m* was detected for Palomero Toluqueño (PAT) and Mushito (MUS), highland Native Maize Varieties.
